# Rastermap: a discovery method for neural population recordings

**DOI:** 10.1101/2023.07.25.550571

**Authors:** Carsen Stringer, Lin Zhong, Atika Syeda, Fengtong Du, Maria Kesa, Marius Pachitariu

## Abstract

Neurophysiology has long progressed through exploratory experiments and chance discoveries. Anecdotes abound of researchers setting up experiments while listening to spikes in real time and observing a pattern of consistent firing when certain stimuli or behaviors happened. With the advent of large-scale recordings, such close observation of data has become harder because high-dimensional spaces are impenetrable to our pattern-finding intuitions. To help ourselves find patterns in neural data, our lab has been openly developing a visualization framework known as “Rastermap” over the past five years. Rastermap takes advantage of a new global optimization algorithm for sorting neural responses along a one-dimensional manifold. Displayed as a raster plot, the sorted neurons show a variety of activity patterns, which can be more easily identified and interpreted. We first benchmark Rastermap on realistic simulations with multiplexed cognitive variables. Then we demonstrate it on recordings of tens of thousands of neurons from mouse visual and sensorimotor cortex during spontaneous, stimulus-evoked and task-evoked epochs, as well as on whole-brain zebrafish recordings, widefield calcium imaging data, population recordings from rat hippocampus and artificial neural networks. Finally, we illustrate high-dimensional scenarios where Rastermap and similar algorithms cannot be used effectively.

## Introduction

High-density electrodes and two-photon calcium imaging have generated an explosion of large-scale neural recordings [1, 2]. Visualizing and analyzing such recordings can be done either directly at the single-cell level [3, 4], or at the population-level using dimensionality-reduction methods [5–8]. However, both of these have caveats. Visualizing neurons one at a time can be difficult because single neurons are often very noisy [9, 10]. Furthermore, single-neuron visualizations cannot show the population-wide coordination of neural firing patterns, which can vary across trials leading to “trial-to-trial” variability [11–13]. On the other hand, dimensionality reduction algorithms can find common patterns of covariation across neurons, allowing further analyses to be restricted to just these reliable modes of activity. However, a relatively large number of components must be used to capture the high-dimensional structure of the neural activity patterns [14–17]. Furthermore, correlations between these components often make it difficult to visually tease apart their independent contributions to the population activity.

Nonlinear dimensionality reduction methods can overcome some of these limitations. For example, manifold discovery algorithms like t-SNE and UMAP embed the firing patterns of neurons into one or two dimensions [18, 19]. Such algorithms can be used, for example, to place neurons with similar firing patterns close to each other. However, these algorithms are typically used to visualize the embedding space, which is a visualization of the relations between neurons, rather a direct visualization of their activity patterns [20]. Furthermore, it can be challenging for these algorithms to maintain both local and global structure on neural data, as their cost functions are not optimized for such data. Methods like t-SNE and UMAP can also suffer from local minima during optimization [21], and it can be difficult to evaluate what constitutes true clustering in the embedding space and what is an artifact of the algorithms [22].

Unlike these existing methods, Rastermap provides a structured visualization of the activity patterns across different groups of neurons, and shows how these activity patterns morph into each other continuously and how they relate to each other globally. The Rastermap visualization is inspired from “classical” population raster plots, where the spike train of each neuron is shown as a row of rasterized ticks [23, 24]. These raster plots can illustrate the average population activity, but do not show what specific subgroups of neurons do. To improve the plots, one can reorder the neurons across the Y-axis of the plot so that nearby neurons have similar activity patterns (Figure S1). This reordering is the main idea and algorithm behind Rastermap.

The reordering can be done with a one-dimensional nonlinear embedding algorithm. In Rastermap, we developed such an algorithm and optimized it for neural data by combining two commonly observed features of neural activity: 1) a power law scaling of eigenvalue variances and 2) sequential firing of neurons. To overcome local minima, we developed a specialized global optimization framework based on ideas from traveling salesman algorithms. We demonstrate that Rastermap outperforms t-SNE, UMAP and other nonlinear dimensionality reduction methods on neural data. The algorithm is also fast: it runs in less than two minutes on datasets with tens of thousands of neurons. Finally, to evaluate and visualize the patterns identified by the algorithm, we use Rastermap outputs to generate a two-dimensional image of the neural activity, which we sometimes refer to as a “Rastermap”. We illustrate Rastermap on a variety of neural datasets from different species (mouse, rat, zebrafish) as well as on artificial neural networks. Rastermap is implemented in python, and can be run in a jupyter notebook, on the command line, or in the provided graphical user interface (Figure S2).

## Results

The goal of Rastermap is to obtain a sorting of all neurons in a recording, such that nearby neurons in the sorted list have similar functional properties. Additionally, we would like the neural pairwise similarity to decay smoothly as a function of how far the neurons are in the sorting. Equipped with this sorting, we can make a single raster plot of all neurons that visualizes the most common patterns of activity. To start, we cluster the neural activity profiles by k-means clustering, typically into *N*_clusters_ = 100 distinct clusters (Figure 1a). We then define an asymmetric similarity measure between clusters, as the peak cross-correlation between the cluster activities at non-negative timelags (Figure 1a,b). The asymmetry induced by this metric ensures that a well-defined ordering can be achieved, so that clusters with earlier activity are typically displayed towards the bottom of the raster plots.

**Figure 1:**
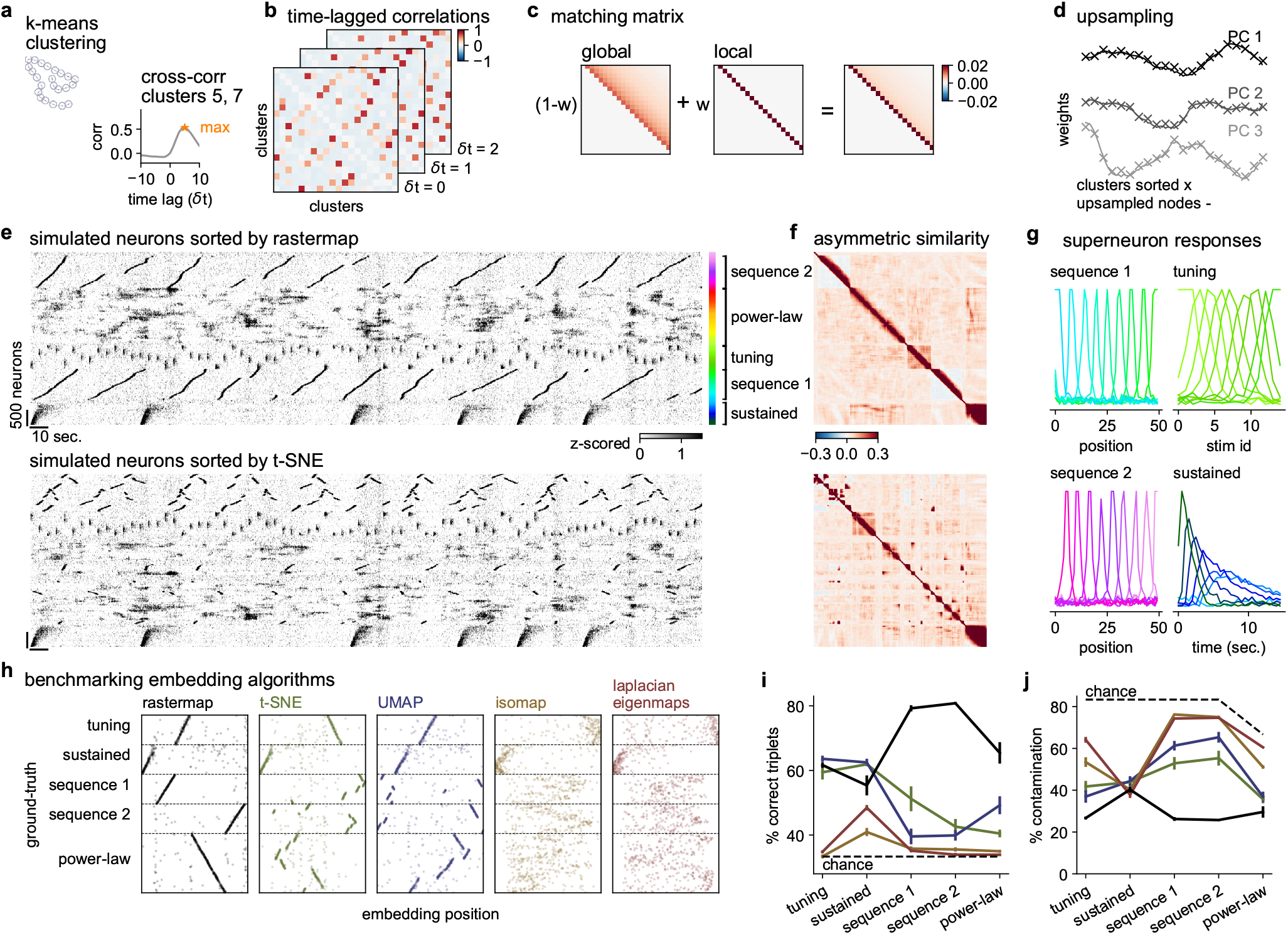
Benchmarking Rastermap on simulated data with multiplexed neural activity. **a-d**, These panels illustrate how Rastermap works. **a**, First, Rastermap divides neurons into 50-200 clusters based on their activity (left). The cross-correlations between different clusters are computed at several time lags (right). **b**, The cluster correlations at different positive time lags are shown for a subset of clusters, and the maximum of these matrices over a time window from 0 to *T*_*max*_ defines an “asymmetric similarity matrix”. **c**, The asymmetric similarity matrix is sorted to match the “matching matrix”, which is a sum of a global similarity matrix and a local similarity matrix. **d**, The cluster features are upsampled using a locally-linear interpolation method, then each neuron is assigned to an upsampled cluster center. **e**, The simulated neurons were sorted by Rastermap or t-SNE, then averaged in bins of 50 neurons – the averages of these neurons are called “superneurons”. **f**, The sorted asymmetric similarity matrix for the simulation. **g**, The activity of the superneurons aligned to different stimulus events. **h**, The sorting of neurons from various algorithms plotted against the ground truth sorting. **i**, For each module of the simulation and each algorithm in **h**, the percentage of correctly ordered triplets is shown. **j**, The percentage of contamination in a module with neurons from other modules.

Having obtained an *N*_clusters_ by *N*_clusters_ similarity matrix, the optimization goal of Rastermap is to permute the rows and columns of this matrix until it matches a predefined matrix as close as possible. The pre-defined matrix is chosen as a sum between a global and a local similarity matrix (Figure 1c, see also multi-perplexity t-SNE [25, 26]). The global similarity decays smoothly as a function of distance between clusters, and the smoothness is chosen so that the eigenvalues of the matrix decay as 1*/n*, which is typical of neural population activity. The local part of the matrix has a “traveling salesman” structure [27], where the similarity is only high between consecutive nodes in the sorting, thus capturing the sequential activity patterns often seen in neural datasets. The local and global matrices are added together typically in equal parts, although the weightings can be adjusted based on the properties of the data. The resulting matching matrix is the target that must be matched to the neural similarity matrix by permuting rows and columns. This optimization is susceptible to local minima, thus we perform it using global optimization techniques. At every iteration, we exhaustively check if any consecutive sequence of *N* clusters can be moved to any other position in the sorting, starting with sequences of length 1 and then progressively checking longer sequences, extending beyond 2- and 3-length sequences which are often used [28, 29]. This global optimization can be implemented efficiently on modern CPUs using the numba python package [30], as long as *N*_clusters_ ≤200. After re-sorting, the neural similarity matrix resembles the matching matrix, as illustrated on an example simulation in Figure 1f. Having obtained an ordering for the clusters, we must now obtain an ordering for the neurons. To do this, we upsampled the sorted cluster activities by a factor of 10 in the PCA feature space, thus creating *N*_clusters_ × 10 positions that can be matched to single neurons (Figure 1d). Single neurons were then assigned to the position that is most highly correlated to their activity in PCA space. When the number of neurons is very large (thousands or more), we cannot visualize them as rows of a Rastermap due to a lack of vertical pixels on most monitors. We therefore bin the thousands of neurons into hundreds of “superneurons”. Superneurons are averages across groups of neurons that were put next to each other in the Rastermap, which by definition have similar firing patterns. An added bonus of creating superneurons is that they have less noisy activity compared to single neurons [31].

### Benchmarking with known ground truth

Benchmarking visualization methods is difficult, since a good visualization should be evaluated based on its ability to simplify complex data and this is hard to measure for real datasets. The approach we take here is to start with a realistic simulation of neural activity, which contains multiple, complex signals with different spatiotemporal signatures. We then randomly shuffle the neurons and ask different methods to undo the shuffling. The simulated populations contain multiple submodules with realistic firing patterns: we use two modules with sequential firing, modeling for example place cells when an animal runs through a linear corridor; we then add a module of sensory responses to repeated flashed stimuli where the neurons have tuning curves to these stimuli; we also add a module of neurons with different response durations and latencies to a single stimulus presented many times; finally we add a module of neurons with power-law PCA structure (Figure 1e), and add small amounts of this module to all other modules, to model the effect of spontaneous, ongoing activity [15].

Rastermap is able to find the natural ordering of this simulation. Other methods fail for different modules, typically oversplitting clusters and positioning the pieces far from each other. For this simulation, the visualization illustrates how Rastermap was better able to find the primary patterns in the data, while other methods like t-SNE would suggest a more disordered organization of neural activity, with many unrelated groups of neurons performing different functions (Figure 1e). After sorting, the asymmetric similarity matrix contains high values closer to the diagonal in Rastermap compared to other methods, like t-SNE (Figure 1f). The superneurons, defined as averages of 50 consecutive neurons in the Rastermap sorting, have clearly defined tuning properties, whether as part of a sequence or in response to the simulated stimuli (Figure 1g).

To benchmark a set of commonly-used embedding algorithms, we compare their embedding order with that of the ground truth (Figure 1h). While there are no relations between modules, within each module we expect a non-interrupted monotonic relation between the embedding position and ground truth. To quantify the similarity of these orderings, we use two measures: the number of correctly ordered triplets and the percent contamination of neuron groups from the same module. The fraction of correctly ordered triplets was higher for Rastermap across the two sequence modules and the power-law module compared to all other methods (Figure 1i). For the flashed stimulus response modules (tuning and sustained), the fraction of correctly ordered triplets was similar between Rastermap, t-SNE and UMAP. Other methods like Isomap and Laplacian eigenmaps lagged behind substantially across all modules. Finding correctly ordered triplets is not sufficient to ensure a good ordering; these triplets also have to be part of a continuous, unbroken module. To estimate how broken-up a module is, we quantified the percent contamination with other modules for the neurons sorted in-between any two neurons from the same module (Figure 1j). This contamination was lowest for Rastermap across the sequence modules, the tuned responses and the power-law module, and similar across algorithms for the sustained module. Overall, Rastermap had much lower contamination compared to all other algorithms. Additionally, we show that Rastermap also performs better on the k-nearest neighbor metric introduced by [25] (Figure S3).

### Rastermap on 50,000 neuron recordings during spontaneous behaviors

To illustrate Rastermap in practice we apply it to a variety of datasets. We start in this section and the next, with datasets collected in our own lab, using twophoton calcium imaging of large neural populations of up to 70,000 simultaneously-recorded neurons at sampling frequencies of ∼3.2 Hz [32–34]. First we apply Rastermap to data collected during spontaneous neural activity, where the animal is head-fixed on top of an air-floating ball in complete darkness, without any explicit task [15, 35]. In this preparation, we used a long “D”-shaped coverslip that covers many different cortical areas, including the anterior part of visual cortex, the sensorimotor cortex and the posterior part of motor cortex (Figure 2a). Neurons across the brain had some degree of spatial clustering, as can be seen by their average position in the Rastermap sorting, indicated by the color of the dots. Over a period of two minutes, the populations of neurons visible in Rastermap engaged in a variety of activation patterns that lasted from *<* 1 sec to tens of seconds, with many of the patterns repeatable within this time window (Figure 2b). Overall, the patterns could be divided into two classes based on whether they were active during running or during sitting [36, 37], but within those classes, different subsets of neurons were active at different times.

**Figure 2:**
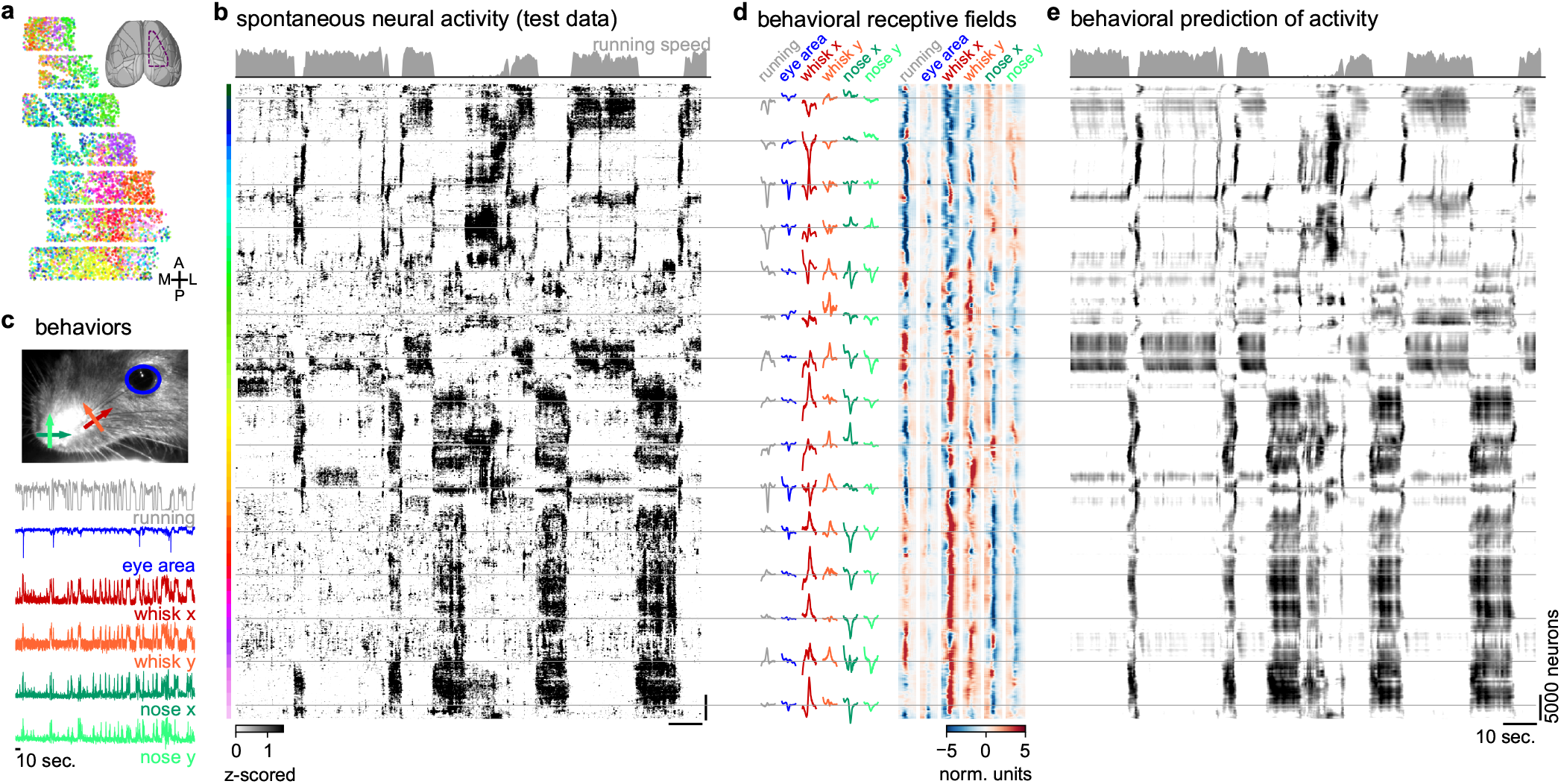
Sorting of spontaneous activity by Rastermap. **a**, 34,086 neurons recorded across mouse sensorimotor cortex using two-photon calcium imaging, colored by position in the Rastermap sorting. **b**, Neural activity sorted by Rastermap. **c**, Mouse orofacial behaviors during the recording. **d**, Left: Example behavioral receptive fields for the superneurons at the Rastermap positions represented by the gray lines. Right: behavioral receptive fields for all superneurons in the Rastermap. **e**, Prediction of neural activity using the behaviors in **c**. The same Rastermap sorting of neurons as in **b** was maintained.

We have previously shown that many of these spontaneous activity patterns can be predicted based on the orofacial behaviors of the mice, which we quantified with either a PCA-based decomposition of the face motion energy or by tracking keypoints on the mouse face [15, 35]. To help us interpret the spontaneous activity clusters, we computed the eye area, whisker position and nose position estimates from the mouse face video using keypoint tracking (Figure 2c) [35]. We then used these behavioral variables to estimate the spatiotemporal linear receptive fields for each superneuron from Rastermap, and also to predict the superneuron activity across time (Figure 2d,e). The receptive field is the spatiotemporal pattern of keypoint movements which would activate that particular superneuron the most. Across the Rastermap embedding dimension, the receptive fields change gradually and appear to be organized hierarchically (Figure 2d). The keypoint with the most influence on superneuron responses was the whisker horizontal location, which separates into negative deflections for the top clusters in the plot (i.e. forward whisker deflections) and positive deflections for the bottom clusters (i.e. backward whisker deflections). Within the set of clusters with negative whisker deflections, a subset were activated positively by running and a subset were activated negatively. Analyzing the patterns of responses on the Rastermap plot itself, we observe different groups of neurons that are activated at the beginning and end of running, and those groups typically were inhibited by running in the model, but excited by whisking. Since whisking typically precedes running and continues briefly after stopping, these patterns of neural responses could be explained as a balanced excitation and inhibition between whisking (excitation) and running (inhibition). This is a novel hypothesis that we generated from analyzing the Rastermap, and it illustrates the kinds of ideas and intuitions that can be inspired by such visualizations. It is however beyond the scope of the present paper to further test this hypothesis.

### Rastermap on 50,000 neuron recordings during a task in virtual reality

Next we applied Rastermap to data collected in visual cortex during navigation and sensory decision-making in virtual reality (Figure 3a) [38–40]. Mice were trained to run through two corridors with different naturalistic textures on the walls (“leaves” and “circles”) (Figure 3b). Reward was delivered at pseudo-random positions in the leaves corridor, following an auditory cue, and the mouse had to lick to trigger the reward. After a few weeks of training, the mouse learned to reliably lick only in response to the cue in the leaves corridor, and not in response to the cue in the circles corridor. The neural activity generated in this task followed clear sequential patterns, which Rastermap was able to group together (Figure 3c,d). Two large populations of neurons can be seen encoding the circles and leaves corridor, with a slightly larger population encoding the rewarding corridor (Figure 3e). In-between these two populations we observe a third population which encodes the gray-space an area between corridors without visual stimuli. The encoding of the gray-space was also sequential as a function of position, and mostly did not depend on either the previous or the next corridor. The sequential activity was interrupted in the leaves corridor at times when the mouse stops to collect the reward. There are also multiple reward-related populations of interest visible in just the single plot from Figure 3. We leave these as a challenge to the reader to identify, and we will describe those populations in detail in a follow-up study about this task. Finally, there are other populations of neurons which do not seem engaged by any aspects of the task, which we believe to be related to the spontaneous orofacial behaviors.

**Figure 3:**
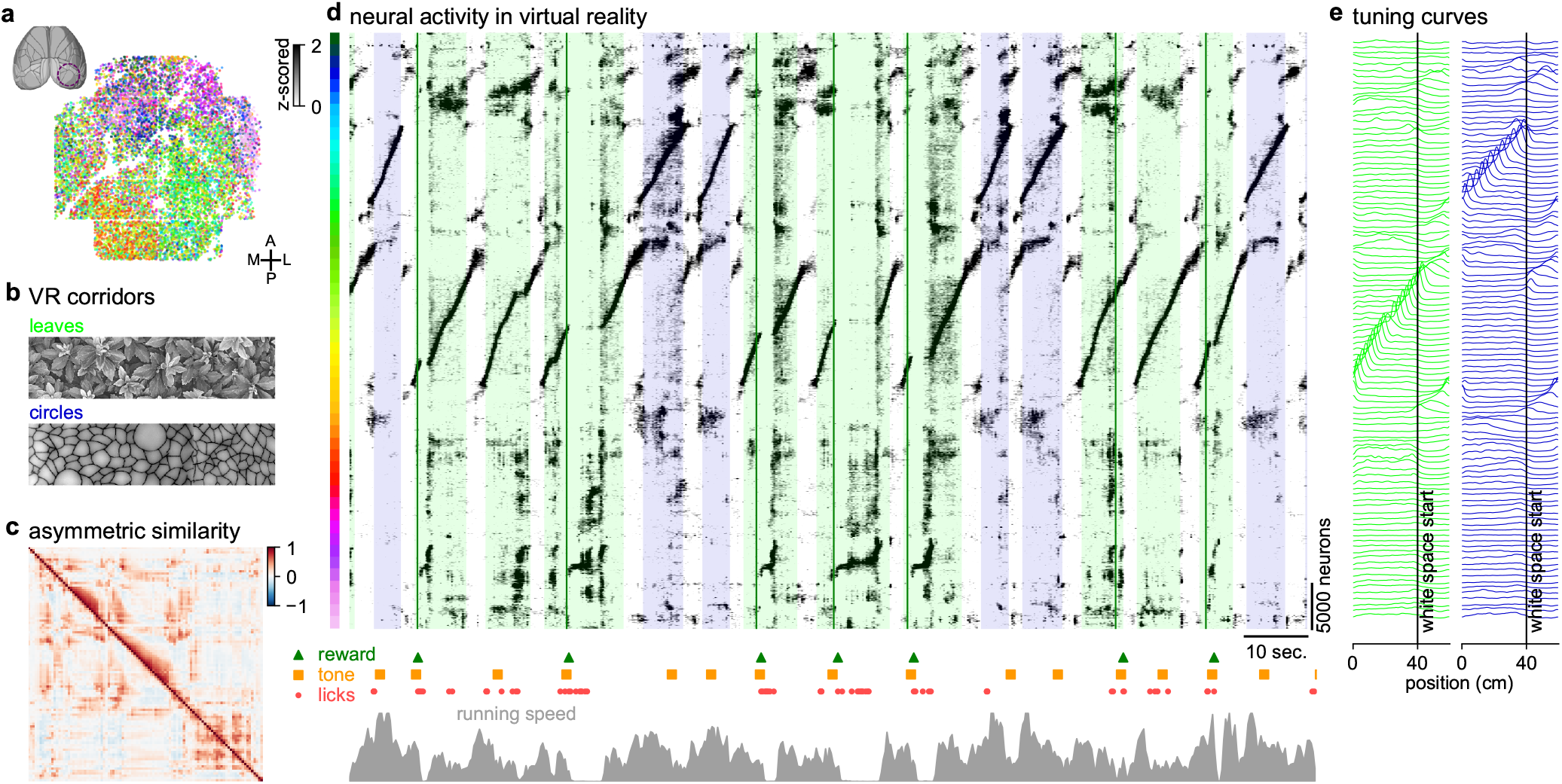
Applying Rastermap to neural activity from a virtual reality task. **a**, 66,318 neurons recorded across mouse visual cortex using two-photon calcium imaging, colored by position in the Rastermap sorting. **b**, During the recording, mice navigated through a 1D virtual reality (VR) with two different corridors (“leaves” and “circles”) which were separated by a gray area and randomly interleaved. A tone was played in each corridor at a random time, and in the “leaves” corridor the tone was followed by a reward. **c**, The sorted asymmetric similarity matrix from the recording. **d**, Top: Neural activity sorted by Rastermap, colored backgrounds denote the type of corridor, green lines denote rewards. Bottom: Event times in the task and running speed. **e**, Superneuron tuning curves to positions along each corridor.

### Rastermap on other biological and artificial neural networks

We have so far illustrated Rastermap on large-scale calcium-imaging data from mouse cortex. In this section, we show that Rastermap can be applied more broadly to recordings from other organisms, with fewer recorded neurons and even on bulk neural activity such as from widefield one-photon imaging. Finally we show an application of Rastermap to artifical neural networks that are used to control agents which play Atari games.

When fewer neurons are recorded (*<*200), Rastermap can skip the k-means clustering step and directly order the neurons according to their asymmetric cross-correlogram peaks. This also allows us to skip the upsampling step, thereby simplifying the algorithm substantially. We applied this simplified version of Rastermap to electrical population recordings from rat hippocampus during running through a linear track (Figure 4a) [41, 42]. Rastermap found two main groups of neurons encoding forward and backward runs through the track. For each group, a subset of neurons encoded the stationary periods at the end of each run. Finally, another group of neurons had dynamics that were only driven by running and not selective to corridor position. This group turned out to be composed entirely of fast-spiking interneurons, which had relatively homogeneous activity.

**Figure 4:**
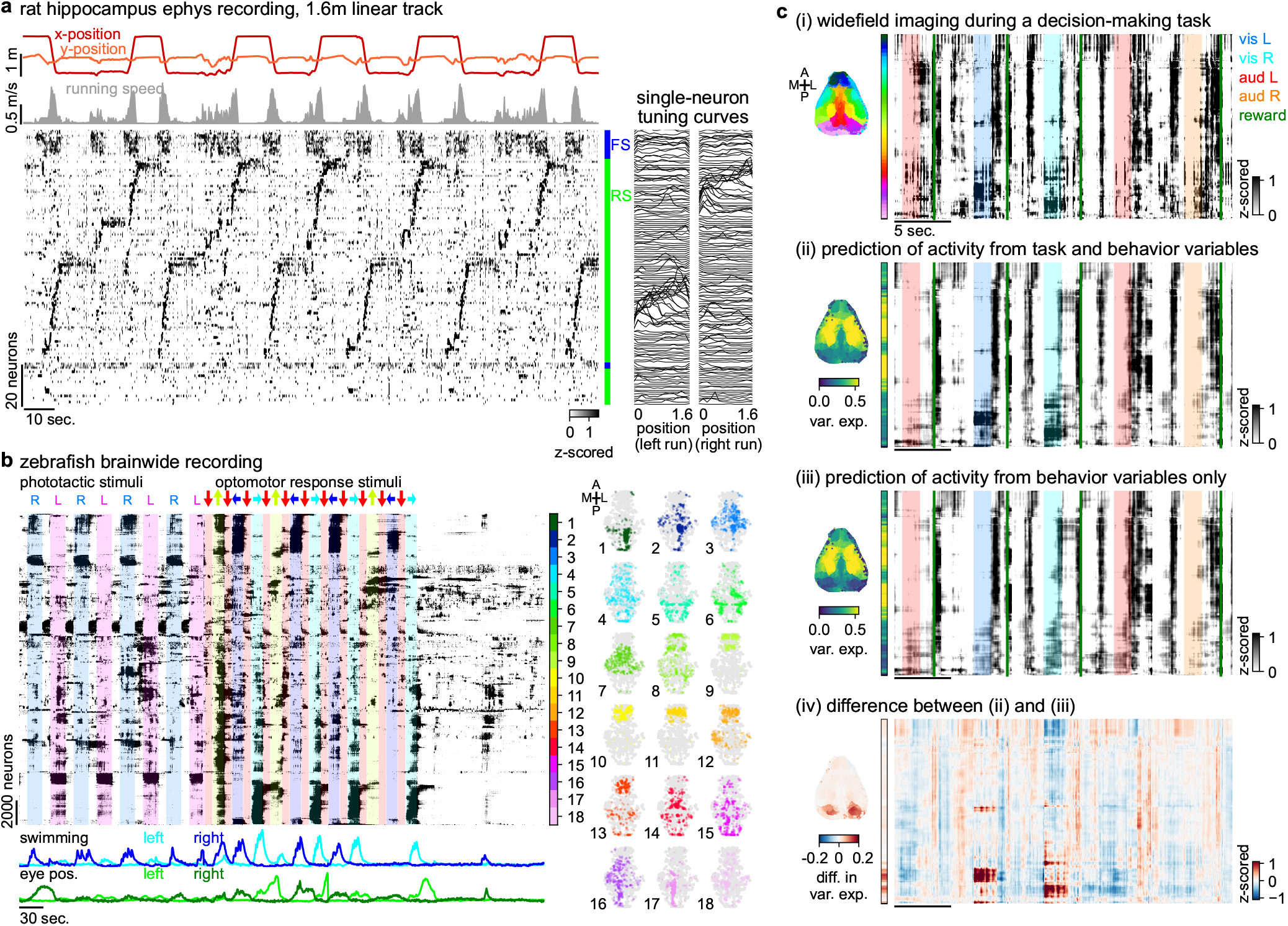
Visualizing neural activity across brain areas and species. **a**, Left: Rastermap sorting of neural activity collected from the CA1 region of rat hippocampus using two multi-shank silicon probes, during which the rat ran back and forth along a 1.6 meter linear track [41, 42]. Neuron identity was defined by spike waveform shape: FS = fast spiking (putative interneuron), RS = regular spiking (putative pyramidal cell). Right: tuning curves of each neuron to the position of the rat in the track. **b**, Left: Rastermap sorting of neural activity collected from the whole-brain of a paralyzed zebrafish using light-sheet imaging at a rate of 2.1Hz [20, 43]. This period of time included two visual stimuli: phototactic stimuli (one side of the screen is dark) and optomotor response stimuli (moving gratings). Middle: Colorbar sectioning Rastermap into 18 groups for visualizing spatial patterning. Right: Neuron positions in each group plotted in color, all neurons in the recording are in gray. Positions in each plot were collapsed across the Z dimension. **c**, (i) Rastermap sorting of cortical activity collected by widefield imaging in mice performing a decision-making task [44, 45]. The voxels in the brain image are colored by Rastermap position. Stimulus events in the task are indicated by colored shaded regions, and reward times are indicated by green lines. (ii) Linear prediction of activity from task and behavior variables shown with same sorting as in (i). (iii) Same as (ii) with linear prediction from behavior variables only. (iv) Difference between prediction in (ii) and (iii).

Another use-case for Rastermap is in multi-area or whole-brain recordings, such as from larval zebrafish using calcium imaging [46]. In this case, different groups of neurons may be identified that correspond to combinations of brain areas that perform a certain function together. We used recordings where different optomotor paradigms were presented using different visual stimulation conditions (Figure 4b) [20, 43]. One condition was during phototactic stimuli, which elicited movement towards bright areas. A second condition was during drifting gratings, which elicited optomotor responses towards the direction of the stimulus, primarily when the stimulus moved left and right. A third condition was under no visual stimulation, in which the fish only rarely swam. Sorting with Rastermap, we found that many activity clusters correlated strongly with swimming in a condition-dependent fashion. Groups of neurons active during visually-evoked swimming typically did not activate during spontaneous activity. However, the groups of neurons active during spontaneous activity typically did have activity during the visual stimulation conditions as well, although this activity was not aligned to the sensory stimulation events [14, 47]. This resembles results that we previously found in rodent visual cortex [15]. These clusters were of two main kinds: 1) spread out throughout the fish brain; 2) concentrated in the anterior, forebrain areas. Another aspect of note were neuron clusters which were active for directional swimming regardless of condition (phototactic or drifting), while other clusters were only tuned to swim direction for specific conditions. These clusters generally aligned to sensory (frontal) and motor (posterior) areas in the fish brain, but substantial regions of overlap existed as well. Similarly, brain lateralization was apparent for most motor-related clusters, but some neuron groups from the other hemisphere were also sometimes included.

Rastermap can also be used on bulk signal recordings, such as from widefield, one-photon calcium imaging in mice [48]. With this method, signals can be recorded from across the entire rodent cortex, but not at single-cell resolution. Instead, each pixel may correspond to the averaged population activity at that location. We used widefield recordings collected while the mouse performed a decision-making task, and during which several behavioral variables were monitored (Figure 4c) [44, 45]. Since different cortical areas can engage for different behaviors, Rastermap can group together brain areas according to the similarity of their dynamics. As expected, the grouping had well-defined spatial relations (Figure 4ci). To start, the embedding was symmetric across hemispheres, with the left and right brain areas embedded at similar locations. Second, the most anterior pixels corresponded to the olfactory bulb and can be seen to have substantially different patterns of activity, which may be linked to sniffing bouts. The more posterior pixels also had different patterns of activity (corresponding to pink and red hues in the plot), and these may have been grouped together by visual responsiveness. To test this hypothesis, we predicted the pixel activities from task and behavior variables, and found that these variables alone can explain activity across the entire Rastermap, including the anterior pixels in olfactory areas (Figure 4cii). We compared this prediction to the prediction using behavior variables alone, which does not include information about stimuli (Figure 4ciii). The main difference between these two predictions were during visual stimulus trials, specifically in the visual cortex (Figure 4civ).

Finally, we also ran Rastermap on artificial neural networks that have been trained with reinforcement learning techniques to play Atari games. We used pretrained networks from DQN (deep Q networks) agents, and clustered all neurons from across all layers of the DQN, illustrating four example games: Pong, SpaceInvaders, Enduro and Seaquest. In all cases, an episode consisted of a single playthrough of the respective game. In games with more repetitive action sequences, like Pong, Rastermap found the repeated neural sequences that corresponded to each repetition, and separated the forward portion of the sequence (ball moving right) from the backward portion (ball moving left). The details of each volley were encoded in the finer details of the neural activity. For games with less structured states, like SpaceInvaders and Seaquest, Rastermap still found sequences of neurons that tend to activate together, but these sequences were more disorganized. In the case of the Enduro game, neural activation patterns were dominated by the graphical context of the game, which changed from day to night and between weather conditions such as “fog” and “ice”. Within each graphical state, mostly non-overlapping groups of neurons were active. A small set of neurons were active in more than one context, and these were generally found in the higher, more “abstract” layers of the deep neural network. In all games, the neurons from the value network were placed all together in the Rastermap, and appeared to have very homogeneous activity that directly corresponded to the value of a state. This indicates that perhaps the value network did not get sufficient gradient information to differentiate the activity of its neurons.

### Space-filling curves for higher intrinsic dimensionality

Rastermap is primarily a visualization algorithm, but visualizations can sometimes be deceptive, especially when the source data is high-dimensional. In this section, we illustrate some use cases where Rastermap is ineffective at finding structure and we try to provide an intuitive understanding of such cases. Take for example Rastermap applied to neural responses in primary visual cortex to a large set of natural images. In this case, the natural images drive very high-dimensional response patterns across cortex, as we previously described [16]. Such data is unlikely to be well described along any one-dimensional embedding dimension, and consequently the Rastermap appears to have a high-dimensional, un-clustered aspect, except for the gain modulation induced by running. The running-modulated clusters correspond primarily to neurons in higher-order visual areas, that have less sensory tuning (Figure S5a,b). Re-sorting the stimuli according to similarity reveals some additional global structure, but the plot is still disorganized (Figure S5c). Computing the linear receptive fields of superneurons from the Rastermap, we see that nearby super-neurons do in fact have similar receptive fields despite their apparently unorganized responses in the Rastermap (Figure S5d). However, these receptive fields cannot be described by a single parameter as would be needed for a good embedding. The filters require several parameters to be well-described: their retinotopic coordinatesr, their orientation, their spatial frequency etc. To arrange filters with these properties across a one-dimensional continuum, Rastermap has to fill up this high-dimensional space with a so-called “space-filling curve”. Similar conclusions can be reached when applying Rastermap to the visual responses of an artificial deep neural network (Figure S5e-g).

To better illustrate when “space-filling curve” behavior occurs, we constructed a simulation where the underlying intrinsic dimension was two, and neurons were described with place field like responses. Sorted with one-dimensional Rastermap or t-SNE, the neurons were arranged across a curve that meandered in a fractal way to fill up the two-dimensional space (Figure S6a). We also constructed a version of Rastermap where the sorting was further broken up iteratively into sub-segments that were sorted with Rastermap again. After three consecutive splits, the simulated population was split into 800 rather than just 100 clusters, and this in turn resulted in a higher-resolution of the underlying space-filling curve. This higher-resolution resulted in better metrics of percent neighbors preserved at small neighborhood sizes, without affecting the number of preserved neighbors at the larger neighborhood sizes (Figure S6b). Thus, while this iterative version of Rastermap can improve on some metrics, it is unlikely to provide fundamentally better visualizations, since the fractal nature of the space-filling curve makes the visualizations un-intuitive (Figure S6c). While a two-dimensional embedding algorithm could be employed, the results of such an algorithm cannot be then used to make raster maps of neural activity.

Our recommendation in these cases is to recognize that Rastermap is fundamentally a dimensionality-reduction method: clustered activity can be found and illustrated in the Rastermap, but higher-dimensional structure may be discarded and missed when it exists. We recommend using other approaches to find and illustrate such high-dimensional structure, such as constrained matrix decomposition techniques (NNMF, seqNMF, ICA, TCA, dPCA, sparse coding), or nonlinear dimensionality reduction with multiple dimensions (t-SNE, UMAP, LFADS, pi-VAE, CEBRA) [5, 51–57].

## Discussion

Here we described Rastermap, a visualization method that can be used to find new, interesting patterns in large-scale neural data. Rastermap makes a two-dimensional plot of neural activity versus time, allowing the user to observe complex spatiotemporal dynamics in relation to experimental events. At the core of the method lies a sorting algorithm which reorders the (possibly) tens of thousands of neurons so that nearby neurons in the sorting have similar activity. The sorting algorithm of Rastermap can also be seen as a one-dimensional embedding method, and has several model features that allow it to accurately embed neural data: 1) modelling the long-tailed decay of pairwise correlations between neurons; 2) modelling sequential activity patterns which are often seen in neural data; 3) using a global optimization algorithm which can avoid local minima. These features allow Rastermap to perform better as an embedding algorithm compared to other methods like t-SNE and UMAP, specifically in the case of one-dimensional embeddings for neural-like datasets.

Using Rastermap, we identified for example different groups of neurons in mouse sensorimotor areas corresponding to whisking at the onset of running and to whisking at the offset of running. We also found diverse patterns of activity associated with corridor positions and reward times in mouse visual areas during a virtual reality task. Rastermap applied to rat hippocampal activity revealed the structure of neural firing along a linear track in putative inhibitory and excitatory neurons. In zebrafish brainwide activity, we observed lateralized and non-lateralized activity patterns associated with different motor and stimulus events. Rastermap sorting of widefield neural imaging provided an unsupervised parcellation of the entire cortex, in part according to the predictability of different regions from different task variables. We found that Rastermap could also be used to discover structure in artificial neural networks such as those trained to play Atari games. Finally, using an extension of Rastermap, we explored datasets with higher intrinsic dimensionality, illustrating the limitations of low-dimensional embedding algorithms when applied to such datasets.

Several studies have already used previous versions of Rastermap to visualize neural activity from flies, zebrafish, mice, and primates [15, 16, 35, 58–67]. We hope that Rastermap will continue to be applied to diverse dataset types and we include a graphical user interface so that users can easily run the algorithm and explore their data. We consider Rastermap to be a good first step in examining neural population activity, such as when a new dataset is first obtained. Rastermap can help users find patterns in data, but to fully demonstrate these patterns, appropriate quantitative analyses must be set up afterwards.

## Acknowledgments

This research was funded by the Howard Hughes Medical Institute at the Janelia Research Campus. We thank the Vivarium staff for animal husbandry, Sarah Lindo and Sal DiLisio for surgery support, Jon Arnold for designing headbars and coverslips, Dan Flickinger for microscopy support, and Tobias Goulet for engineering support.

## Author contributions

C.S. and M.P. designed the study and the algorithm. L.Z. and M.P. performed data collection. C.S., L.Z., A.S., F.D., M.K. and M.P. performed data analysis. C.S. and M.P. wrote the manuscript.

## Code availability

Rastermap was used to perform all analyses in the paper, the code and GUI are available at https://www.github.com/mouseland/rastermap. Scripts for running all the analyses in the paper are available at https://github.com/MouseLand/rastermap/tree/main/paper.

## Data availability

All data will be made available upon publication of the study. Previously shared datasets were also used in the study [42, 43, 45].

## Methods

All experimental procedures were conducted according to IACUC at HHMI Janelia.

### Data acquisition

#### Animals

All experimental procedures were conducted according to IACUC. We performed 3 recordings in 3 mice bred to express GCaMP6s in excitatory neurons: TetO-GCaMP6s x Emx1-IRESCre mice (available as RRID:IMSR_JAX:024742 and RRID:IMSR_JAX:005628). These mice were male and female, and ranged from 2 to 12 months of age. Mice were housed in reverse light cycle, and were pair-housed with their siblings before and after surgery.

#### Surgical procedures

Surgeries were performed in adult mice (P35–P125) following procedures outlined in [31]. In brief, mice were anesthetized with Isoflurane while a craniotomy was performed. Marcaine (no more than 8 mg/kg) was injected subcutaneously beneath the incision area, and warmed fluids + 5% dextrose and Buprenorphine 0.1 mg/kg (systemic analgesic) were administered subcutaneously along with Dexamethasone 2 mg/kg via intramuscular route. For the visual cortical windows, measurements were taken to determine bregma-lambda distance and location of a 4 mm circular window over V1 Cortex, as far lateral and caudal as possible without compromising the stability of the implant. A 4+5 mm double window was placed into the craniotomy so that the 4mm window replaced the previously removed bone piece and the 5mm window lay over the edge of the bone. The sensorimotor window was also a double window and it was placed as medial and frontal as possible. The outer window was 7mm by 4.5mm and the inner window was around 1mm smaller in all dimensions. After surgery, Ketoprofen 5mg/kg was administered subcutaneously and the animal allowed to recover on heat. The mice were monitored for pain or distress and Ketoprofen 5mg/kg was administered for 2 days following surgery.

#### Imaging acquisition

We used a custom-built 2-photon mesoscope [32] to record neural activity, and ScanImage [68] for data acquisition. We used a custom online Z-correction module (now in ScanImage), to correct for Z and XY drift online during the recording. As described in [31], for the visual area recordings, we used an upgrade of the mesoscope that allowed us to approximately double the number of recorded neurons using temporal multiplexing [33].

The mice were free to run on an air-floating ball. Mice were acclimatized to running on the ball for several sessions before imaging, and one mouse was trained on a virtual reality task for two weeks prior to the recording. The field of view was selected such that large numbers of neurons could be observed, with clear calcium transients.

#### Visual stimuli

We showed natural images or virtual reality corridors to the mice on three perpendicular LED tablet screens surrounding the mouse (covering 180 degrees of the visual field of view of the mouse). To present the stimuli, we used PsychToolbox-3 in MATLAB [69]. The flashed visual stimuli were presented for 313 ms, alternating with a gray-screen inter-stimulus interval lasting 313 ms. Occasionally, the screen was left blank (gray screen) for a few seconds. The virtual reality corridors were each 4 meters long, and the mouse moved forward in the virtual reality by running.

#### Videography

The camera setup was similar to the setup in [15]. A Thorlabs M850L3 - 850 nm infrared LED was pointed at the face of the mouse to enable infrared video acquisition in darkness. The videos were acquired at 50Hz using FLIR cameras with a zoom lens and an infrared filter (850nm, 50nm cutoff). The wavelength of 850nm was chosen to avoid the 970nm wavelength of the 2-photon laser, while remaining outside the visual detection range of the mice [70, 71].

#### Processing of calcium imaging data

Calcium imaging data was processed using the Suite2p toolbox [34], available at www.github.com/MouseLand/suite2p. Suite2p performs motion correction, ROI detection, cell classification, neuropil correction, and spike deconvolution as described elsewhere [15]. For non-negative deconvolution, we used a timescale of decay of 1.25 seconds [72, 73].

### Rastermap algorithm and implementation

The Rastermap algorithm is implemented in Python 3 using the numpy, scipy, numba and scikit-learn packages, all of which are easy to install on Windows, Linux and Mac operating systems [30, 74–76]. The graphical user interface is implemented using PyQt5 and pyqtgraph [77, 78]. To perform analyses and create the figures in the paper we used jupyter notebooks and matplotlib [79, 80].

The algorithm involves 5 main steps: dimensionality reduction, clustering, computing the asymmetric similarity matrix, sorting this matrix, and upsampling the cluster centers. See Figure 1a-d for a graphical representation of these steps.

#### Data normalization and dimensionality reduction

First we normalize the neural activity in order to avoid fitting single neuron statistics with the embedding algorithm. We z-score the activity of each neuron so that the mean activity of each neuron is 0 and its standard deviation is 1. Then we project out the mean across neurons at each time point, this is optional but recommended (parameter `mean_time` in the algorithm). An optional step, which depends on the noise level of the data, is binning in time, parameter `time_bin`. This binned data is then used to compute the singular vectors.

Next, to make the data size more manageable, we compute the singular value decomposition of this normalized activity matrix, where the left singular vectors will be of length number of neurons. We generally keep the top 100 to 400 left singular vectors, this can be specified by the user with the parameter `n_PCs` in the algorithm. We scale each of these singular vectors by its singular value, to preserve distances in the original space, and compute clusters from these scaled singular vectors.

#### Clustering

We identified groups of coactive neurons using scaled k-means clustering [34]. Compared to regular k-means, scaled k-means fits an additional variable λ_*i*_ for each neuron *i* such that

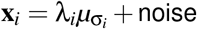

where **x**_*i*_ is the activity vector of neuron *i*, σ_*i*_ is the cluster assigned to neuron *i* and *μ*_*j*_ is the activity of cluster *j*. Like regular k-means, this model is optimized by iteratively assigning each neuron to the cluster which best explains its activity, and then re-estimating cluster means. The number of clusters *N* computed is called the `n_clusters` parameter in the algorithm.

#### Asymmetric similarity matrix

For each cluster out of *N* clusters, we compute the mean cluster activity by averaging all of the neurons in a cluster, and then we z-score each cluster activity trace. We then compute the cross-correlation between all cluster activity traces *c*_*i*_:

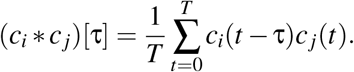

This is computed for a specified number of positive τ time lags τ_max_, called the `time_lag_window` parameter in the algorithm. Then we use the maximum value of the cross-correlation for each cluster pair over these positive τ values for our asymmetric similarity matrix *S*:

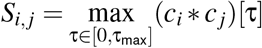

#### Sorting the similarity matrix

We optimize the sorting of the asymmetric similarity matrix of the cluster nodes to maximize a matching score. This score is defined as the dot product between the sorted version of the asymmetric similarity matrix *S*^sorted^ and a pre-specified matching matrix *M*:

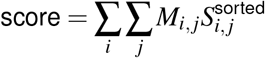

This matching matrix has two parts: a global similarity and a local “traveling-salesman” similarity. The global similarity matrix has eigenvalues which decay with a power law of coefficient 1, defined as

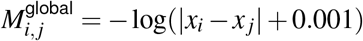

where *x*_*i*_ = *i/N* and the diagonal 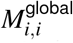 is set to zero. The local similarity matrix is close to 1 on the first off diagonal and very small elsewhere:

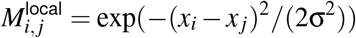

where σ = *i/*(2*N*) and the diagonal 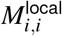 is set to zero. We then set the lower diagonal of each matrix to zero to force all correlations to be put above the diagonal, which enforces forward sequences of activity. Each of these matrices are then normalized by their sums across all entries. Then the final matching matrix is a weighted sum of the two matrices:

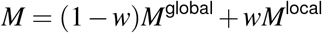

where the weighting *w* is called the `locality` parameter in the algorithm.

We initialize the sorting by the first singular vector weights for each cluster node. Then we compute the change in score for each cluster moved to each position in the matrix (n*(n-1) tested moves). We first test all movements of groups of a single node, and then we move the node that increases the score the most. If none of the moves will increase the score, then we test all moves of two consecutive nodes, and similarly if none of those moves, we test all groups and moves of three nodes, and so on. We repeat each step of searching for moves of groups of nodes that increase the score for 400 iterations, or until no move of any group of nodes increases the score. During this optimization, we skip every other node if the number of clusters to sort is 100 (the number of nodes skipped is the floor of the number of clusters divided by 30). If nodes were skipped, then the optimization is run again with all nodes for up to 400 iterations, although it usually takes fewer than ten iterations to converge during this second run, and thus skipping nodes reduces runtime. This optimization can be made highly parallel: we can test all moves of groups of nodes of a certain length simultaneously, then choose the best move. Therefore we accelerated this step and other steps in the optimization using the numba library [30] in python, which can be easily installed on any standard desktop or laptop computer.

This optimization takes less than 10 seconds for 100 clusters, around 20-30 seconds for 150 clusters, and 1-2 minutes for 200 clusters. Because it exponentially gets slower for larger numbers of clusters, we find more positions for neurons by upsampling rather than using more clusters.

#### Upsampling and superneuron computation

We upsampled in-between cluster centers in PCA space using weighted, locally-linear regression, to go from *N*_clusters_ to 10 · *N*_clusters_ nodes. The regression approximated linearly the function from discrete cluster index to PCA features, in small local neighborhoods around each cluster index. The sizes of these neighborhoods were controlled by weighting the regressors according to their index separation in the Rastermap sorting. The weightings were Gaussian as a function of Euclidean distance with standard deviation 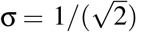. Cluster centers beyond the 50 nearest neighbors of an upsampled point were not used. We performed this linear approximation at each upsampled position, at a resolution of 10x the original resolution of the data. The upsampled features were then correlated with each neuron activity, and neurons were assigned to the position of their best-matching upsampled node.

#### Clustering and splitting steps

To increase the number of clusters, we also explored a strategy inspired by space-filling curves [81]. Starting with 100 sorted clusters, we divide the sorting into quartiles of 25 clusters each, and reclustered the neurons in each quartile into 50 clusters. We then sort each of these groups of 50 clusters with the iterative optimization, including the asymmetric similarity matrix surrounding each group in the score to avoid discontinuities in the final sorted matrix across groups of clusters. The splitting and reclustering can be performed as many times as preferred, the parameter in the algorithm is `n_splits`. For the analyses in Figure S6 and Figure S5, we split three times resulting in 800 total clusters.

### Simulations

In order to compare the performance of different embedding algorithms, we created simulations of largescale data with noise (Figure 1 and Figure S6).

#### Simulation with different modules

We created a simulation with five different types of one-dimensional modules: two sequence modules, one module with tuning curves, one module with sustained stimulus responses, and one module with power law eigenvalues (Figure 1e). The first four modules had 1,000 neurons each and the last module had 2,000 neurons.

Each neuron in a sequence module was assigned to a random position along the sequence at which point it activated. The sequences in the sequence modules repeated many times throughout the simulation, with each repetition having a random length between 350 and 700 timepoints and the time between repetitions having a ranodm length between 100 and 200 timepoints. A variable velocity for each sequence repetition was generated by adding to a constant velocity random gaussian noise filtered by a Gaussian with standard deviation 30 timepoints. These sequences could break at a random place for a short period with a probability of 50% per repetition, simulating the breaks in sequences we observed when mice stopped moving through virtual corridors.

The tuning curve module consisted of neurons with one-dimensional Gaussian tuning curves at 1500 possible positions along this axis. We presented in a random order 15 stimuli equally-spaced along this one-dimensional axis. In total there were 500 stimulus presentations spaced 100 timepoints apart. The stimulus responses from the neurons decayed exponentially from the onset with a time scale of 25 timepoints.

The sustained module consisted of neurons with varying latencies and response durations to a single stimulus generated using a difference of exponentials filter with 100 timescales ranging from (25, 5) to (304, 61). Each neuron in the modules was assigned randomly to one of these 100 timescales. The inter-stimulus interval was drawn from an exponential with a decay time scale of 750 timepoints, with a minimum value of 2,000 timepoints used.

The power law module consisted of a neural population with singular values *s*_*k*_ = 1*/k*^2^. The right singular vectors of the neural population *V*, with length time, consisted of a sparse boolean matrix filtered by an exponential filter with a time scale of 25 timepoints. The left singular vectors *U* were composed of cosine functions with increasing frequency as a function of component number *k*: cos(π*kx*), where *x* is a random number for each neuron between 0 and 1. Each neuron in the entire simulation is assigned a random *x* value. *U, S*, and *V* were then multiplied and clipped at zero to create a positive neural activity matrix. We then added this activity to every neuron in the simulation; this reproduced the property in the data that a majority of neurons are driven during spontaneous activity periods, and they continue to be driven by spontaneous activity patterns during stimulus presentations. 2,000 neurons out of 6,000 were driven solely by the power law module.

The activity matrices created in each module represent the firing rate of each neuron at each timepoint. We scaled each firing rate trace by a random number drawn from an exponential to create neurons with different firing rates. We then used the Poisson distribution to generate spikes from these firing rates. We also added independent Poisson noise to all the neurons with mean 0.03.

#### Simulation with intrinsic dimensionality of 2

We simulated neural activity with an intrinsic dimensionality of 2 by randomly choosing an *x* and *y* value for each neuron in the range of 0 to 1 (Figure S6). We simulated 30,000 neurons in total using basis functions which depended on *x* and *y*: 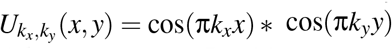, where and *k*_*x*_ and *k*_*y*_ compose a 2D grid from 1 to 30, for a total of 900 basis functions, which we used as the left singular vectors for construction of the simulation. The singular values were defined as 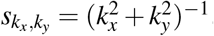. The right singular vectors *V* were generated as random Gaussian noise at each of the 20,000 timepoints. We multiplied *U, S* and *V* and then added random Gaussian noise scaled by 5e-3 at each timepoint.

### Benchmarking embedding algorithms

Each module in the one-dimensional simulation has a ground truth sorting defined by its one-dimensional axis. We compared the embedding order found by a given algorithm to this ground truth order using two metrics: the percent of correctly ordered triplets and the percent contamination of neuron groups from the same module. To compute the triplet score, we drew many groups of three random neurons from the same module and if these three neurons were in the same order in the embedding as in the ground truth then it was considered a correct triplet. For the percent contamination we drew many groups of two random neurons from the same module and quantified the percentage of neurons between these two neurons which were from a different module, and averaged the percentage over all groups. The results are shown in Figure 1i,j.

Another way to benchmark embedding algorithms is to quantify how well the local neighborhood of a data point in the original space is preserved in the embedding space [25]. This is done by computing the percentage of *k*-nearest neighbors in the original space are preserved as *k*-nearest neighbors in the embedding space. In our case, for the original space we can use the ground truth position of the neuron in the simulation and compute distances between these positions, rather than using noisy estimates of distances from the data as is required when the ground truth is unknown. We computed the percentage of neighbors preserved for *k* from 1 to 500 on a random subset of 2,000 neurons from the full simulation (to speed up the neighbor computation); the results are shown in Figure S3.

#### Running other embedding algorithms

We compared Rastermap to the most commonly used embedding algorithsm: t-SNE [18], UMAP [19], ISOMAP [82], and Laplacian Eigenmaps [83] (Figure 1h-j, Figure S3a). We ran the OpenTSNE implementation of t-SNE due to its efficiency and flexibility [84]. We ran t-SNE and UMAP with the suggested initialization from [25]: the first principal component scaled by a small number. We used the cosine similarity metric for t-SNE, UMAP and ISOMAP as this improved performance. Otherwise the algorithms were run with their default parameters.

The performance of t-SNE and UMAP can depend on their parameters which define their local neighborhoods, perplexity and n_neighbors. Therefore, we also ran t-SNE and UMAP with several different values of these parameters to determine whether it influenced the embedding quality (Figure S3b,c).

### Data analysis

#### Spontaneous activity in sensorimotor areas

We analyzed neural activity collected from a large part of mouse dorsal cortex using two-photon calcium imaging at a rate of 3.2Hz, centered on sensorimotor areas, while the mouse was free to run on an air-floating ball in total darkness (Figure 2a). We sorted the neural activity with Rastermap using n_clusters=100, n_PCs=128, locality=0.75, and time_lag_window=5, and binned the sorted neural activity into superneurons of size 50 neurons each (Figure 2b); the neurons are colored by the sorting in Figure 2a.

We used the keypoint tracking network from [35] in order to track keypoints on the mouse face from the video taken during the recording (Figure 2c). From these keypoints we computed five interpretable variables: the eye area, the whisker pad position, and the nose position. The eye area was computed by taking the difference of the top and bottom eye key-points and the difference of the left and right eye key-points, and then multiplying these two values together. The whisker pad position was computed by averaging the positions of the three tracked whisker key-points. Then the principal components of the x- and y-positions were computed and used to rotate the coordinates such that the new x-position corresponded to movements along the major axis of whisker movements. The nose position was computed by averaging the position of the four tracked nose keypoints.

Next we used the neural network from [35] to predict 128 neural activity principal components from these five variables and the running speed. The behavioral prediction from this non-linear neural network was visualized in Figure 2e. To estimate the superneuron receptive fields we used a simplified linear version of this neural network. The network consisted of the same first two layers, an input linear layer and a one-dimensional convolutional layer, and then these layers were followed by a single output linear layer, which predicted the 128 PCs (Figure 2d). The receptive field for a superneuron was estimated by optimizing a small behavioral snippet of length 8 seconds to maximally activate the superneuron at the timepoint at the midpoint of the snippet.

#### Neural activity from a virtual reality task

We analyzed neural activity collected from mouse visual cortical areas using two-photon calcium imaging at a rate of 3.2Hz, while the mouse was free to run on an air-floating ball in a virtual reality task (Figure 3a). The task contained two virtual corridors, “leaves” and “circles”, and the “leaves” corridor was rewarded at a random position in the corridor after a sound cue (the sound cue was also played in the “circles” corridor but not rewarded) (Figure 3b). We sorted the neural activity with Rastermap using n_clusters=100, n_PCs=200, locality=0.75, and time_lag_window=10, and binned the sorted neural activity into superneurons of size 100 neurons each (Figure 3d); the neurons are colored by the sorting in Figure 3a, and the asymmetric similarity matrix for the clusters is shown in Figure 3c. The superneuron tuning curves were computed for 100 positions along each corridor and in the gray-space between corridors.

#### Rat hippocampus data

We analyzed a freely available neural activity recording collected from the CA1 region of rat hippocampus using two multi-shank silicon probes, during which the rat ran back and forth along a 1.6 meter linear track (Figure 4a) [41, 42]. We binned the spiking in time bins of 200 ms, and used the full time period in which the rat was in the maze, and used all 137 recorded neurons. To estimate the location of the rat and the start and stop of the rat in each corridor, we used code available from [56].

We sorted the neural activity with Rastermap using n_clusters=None, n_PCs=64, locality=0.1, and time_lag_window=0. When n_clusters is set to None or to 0, then the algorithm sorts the original datapoints – the single neuron traces – rather than first clustering the data and then sorting. Tuning curves for leftward runs and rightward runs along the corridor were computed for 30 positions along the track.

#### Zebrafish whole-brain data

We analyzed a freely available neural activity recording collected from the whole-brain of a paralyzed zebrafish using light-sheet imaging at a rate of 2.1Hz (Figure 4b) [20, 43]. During the imaging session, the zebrafish was presented many different visual stimuli, such as phototactic stimuli (one side of the screen is dark) and optomotor response stimuli (moving gratings). The fictive swimming was recorded with electrodes and the eye positions tracked. We removed neurons which had low SNR, using a threshold of 0.008 on the fluorescence standard deviation. To remove long timescales from the calcium sensor in the data, we baselined the fluorescence traces and ran non-negative spike deconvolution with a timescale of 2 seconds [72, 73].

We sorted the neural activity with Rastermap using n_clusters=100, n_PCs=200, locality=0.1, and time_lag_window=5, and binned the sorted neural activity into superneurons of size 50 neurons. We then divided the plot into 18 bins to color neurons across the fish brain by position (Figure 4b right).

#### Widefield imaging data

We analyzed a freely available widefield cortical imaging recording collected from mice performing a decision-making task (Figure 4c) [44, 45]. We discared voxels on the edges of the recording area as these voxels were noisy in time, but this step is optional and data-dependent. 186,590 voxels remained for analysis, by 93,177 timepoints – the data was collected at a rate of 30Hz. Since widefield imaging recordings are very large (hundreds of thousands of voxels by hundreds of thousands of timepoints), they are often summarized by their singular value decomposition. Rastermap has the option to run on these singular vectors alone, rather than the full dataset. We sorted the voxel singular vectors with Rastermap using n_clusters=100, n_PCs=200, locality=0.5, and time_lag_window=10, and binned the sorted voxels into supervoxels of size 200 voxels.

Next, we predicted the supervoxel activity from behavior variables or both behavior and task variables. These were precomputed in [45] in “regData.mat”. The behaviors used were handle grabbing movements, licking, whisking, nose movements, filtered pupil area, face movements, body movements, and principal components of the raw video of the mouse and the video motion energy. The task variables used were reward times, choice, previous choice, water delivery, piezo, visual stimuli, and auditory stimuli. We z-scored each of these variables across time. We predicted the voxel activity from these variables using linear regression with a regularization constant of 1e4. The prediction from behavior-only is shown in Figure 4cii, and the prediction from behavior and task variables is shown in Figure 4ciii. The difference between the two predictions is shown in Figure 4civ.

#### Visual stimulus responses

We analyzed neural activity collected from a subset of mouse visual cortical areas using two-photon calcium imaging at a rate of 3.2Hz, while the mouse was free to run on an air-floating ball and grayscale natural images were presented (Figure S5a). A natural images was shown on every other neural frame, there were 5,000 different images in total, presented 3 times in a random order. To compute linear receptive fields, we downsampled the natural images to size 24 by 96 then computed the top 200 principal components.

We sorted the neural activity with Rastermap using n_clusters=100, n_PCs=400, n_splits=3, locality=0.0, and time_lag_window=0 (resulting in 800 clusters due to splitting), and then binned the sorted neural activity into superneurons of size 139 neurons to create 500 superneurons in total (Figure S5b). We then averaged the responses of each superneuron over the 3 repeats of the 5,000 images. Using the averaged responses, we computed the linear receptive fields of each superneuron with linear regression from the image principal components with a regularization constant of 1e4 (Figure S5c).

Rastermap can also be applied along the time or stimulus axis of the neural activity matrix to examine relationships among timepoints or stimuli. We applied Rastermap along the stimulus axis of the superneuron activity averaged over 3 repeats of the 5,000 images, using n_clusters=100, n_PCs=400, n_splits=0, locality=0.0 (Figure S5c).

### Neural network experiments

#### Deep Q-network playing Atari games

We analyzed the activations of neural networks trained to play Atari games, from the Stable-Baselines3 RL Zoo (Figure 5) [49, 50]. These networks were quantile regression deep Q-networks (QR-DQNs), which consisted of 3 convolutional layers and a linear layer to process the images from the game (4 frames stacked in time) and a feedforward network to compute the state values [85, 86]. We used 4 different “NoFrameskip-v4” agents each trained on a different environment: Pong, SpaceInvaders, Enduro, and Seaquest. We ran the environments ten times, each time with a different random seed, and then concatenated the activations across the ten episodes. Each episode lasted for up to 4,000 timepoints, or however long until the agent won or lost in each run (only for the Enduro environment did this exceed 4,000 timepoints because the Enduro game can never be won, and the agent never lost).

**Figure 5:**
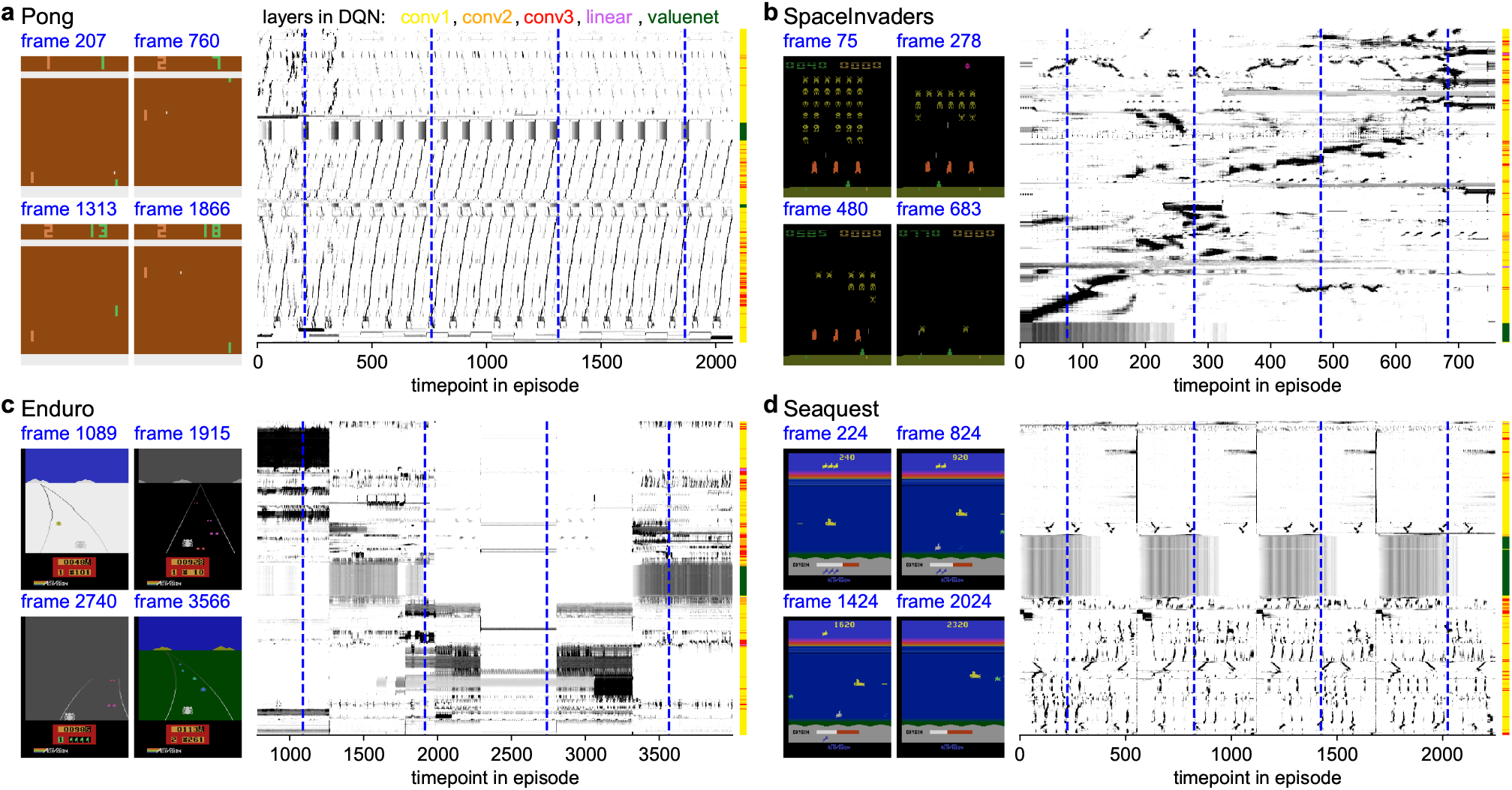
Applying Rastermap to artificial neural networks. We visualized activations from QR-DQN networks that were trained to play various Atari games [49, 50]. **a**, An agent trained on the Pong Atari game. Left: Example frames from an episode of the agent playing Pong. Middle: Activations of units from the agent’s neural network during the episode, sorted by Rastermap. Right: Positions in the sorting colored by the layer in the network. **b-d**, Same as **a**, for agents trained to play Space Invaders, Enduro and Seaquest.

We sorted the neural network activations across all ten episodes with Rastermap using n_clusters=100, n_PCs=200, locality=0.75, and time_lag_window=10, and binned the sorted activations into superneurons of size 50 units. We showed the activations for one episode along with four example frames in Figure 5.

#### Alexnet in response to natural images

We trained the Alexnet neural network to perform image recognition on ImageNet images in grayscale (rather than the usual RGB) [87, 88]. We then presented the network the same natural images that we showed to the mice and saved the activations in all of the layers. For further analysis we used 2,560 random activations from the first four convolutional layers, and all 1,280 activations from the fifth convolutional layer.

We sorted the Alexnet activations with Rastermap using n_clusters=100, n_PCs=400, n_splits=3, locality=0.0, and time_lag_window=0 (resulting in 800 clusters due to splitting), and then binned the sorted activations into superneurons of size 24 units (Figure S5c, right). We colored each of the activations by their position in the rastermap (Figure S5c, left). We applied Rastermap along the stimulus axis of the superneuron responses obtained above using n_clusters=100, n_PCs=400, n_splits=0, locality=0.0 (Figure S5f).

---

**S1:**
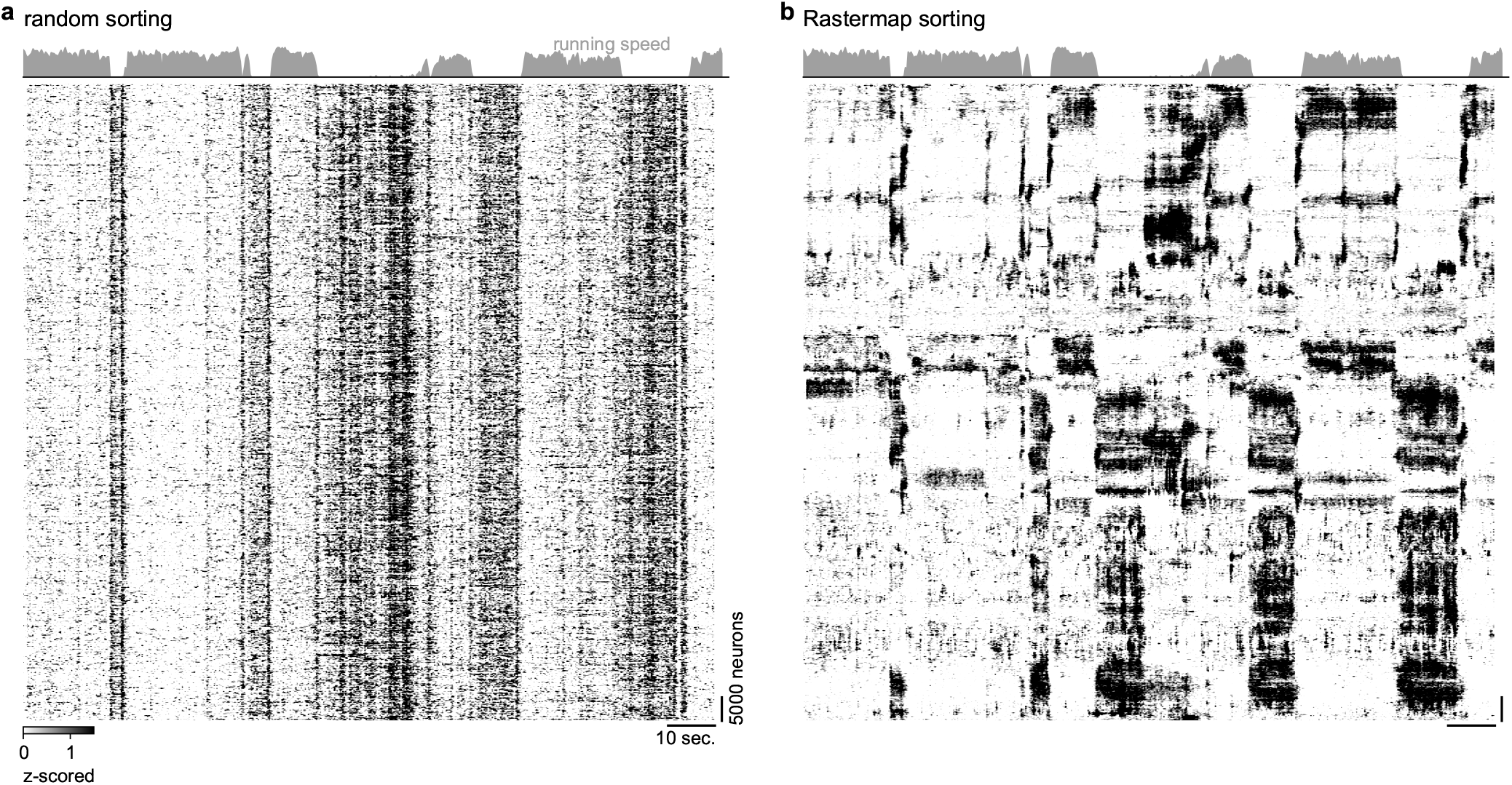
Spontaneous neural activity in random order and sorted by Rastermap. **a**, Same neural activity as Figure2, with random sorting. **b**, Figure2b reproduced for comparison.

**S2:**
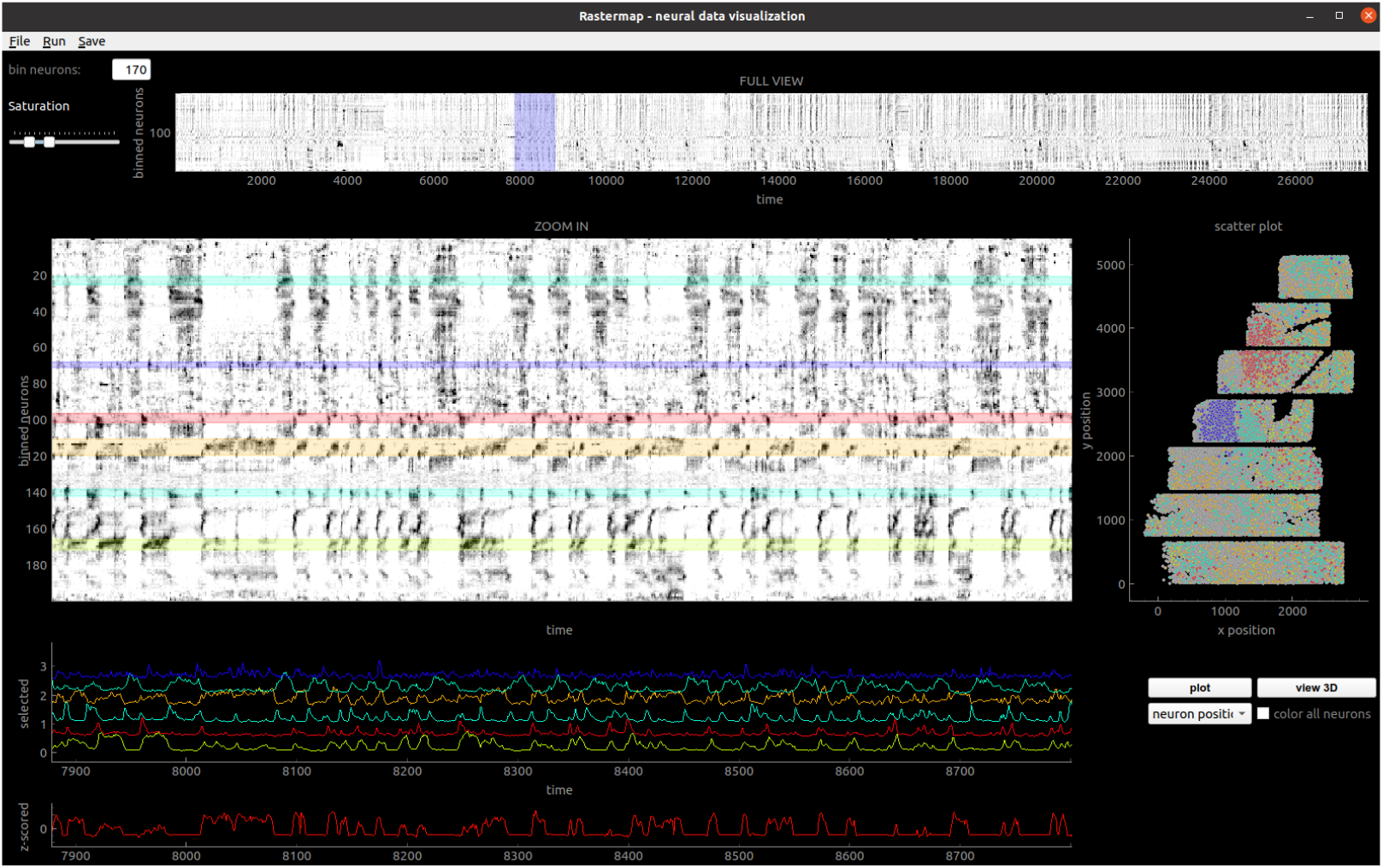
Rastermap graphical user interface (GUI). 1st row: Full recording, with selected time period shown in blue. 2nd row left: Superneuron activity by time in selected time period, with user-selected clusters highlighted in different colors. 2nd row right: All neurons shown in gray, and neurons from selected clusters shown in color. 3rd row: Mean activity in each of the user-selected clusters. 4th row: Behavioral/task variable plotting area showing the mouse’s running speed.

**S3:**
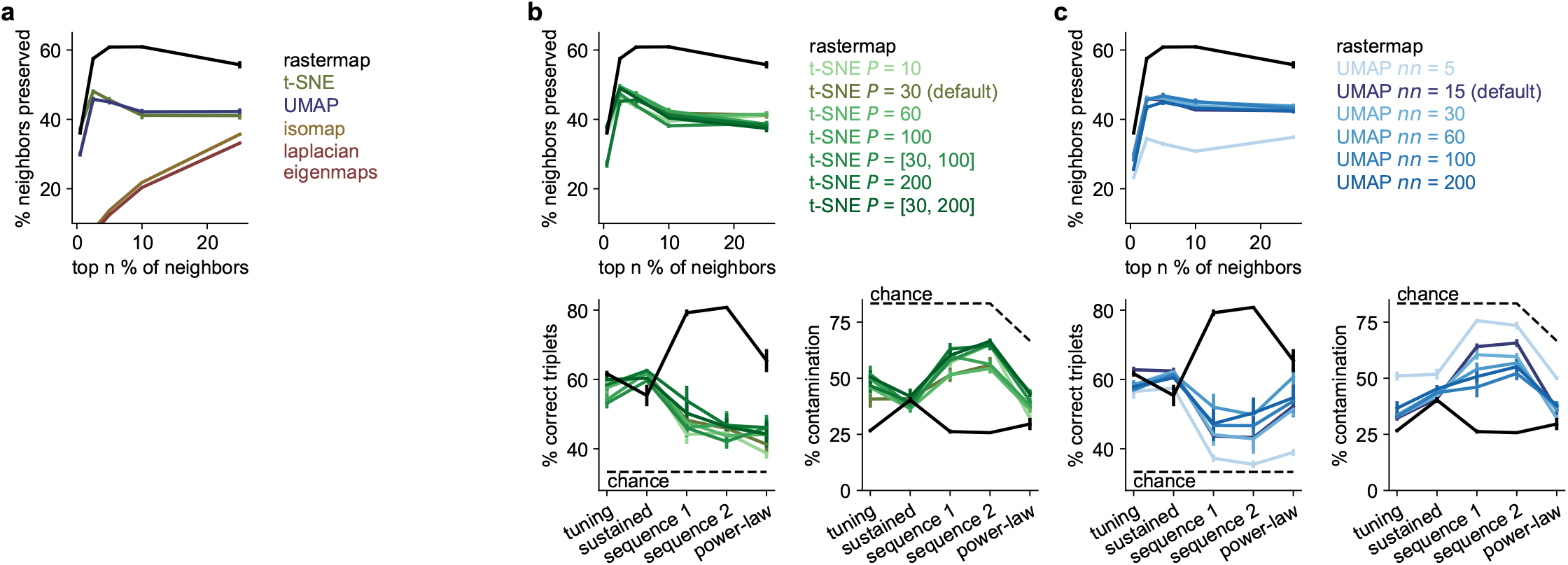
Benchmarking embedding algorithms. **a**, The KNN score for benchmarking embedding algorithms from [25]: the percentage of *k*-nearest neighbors in the original space that are preserved as *k*-nearest neighbors in the embedding space. **b**, Top: Same as **a** for t-SNE embeddings computed with various perplexities, including multiple perplexities [84]. Bottom: Like in Figure1ij, percentage of correct triplets and percentage contamination for the different t-SNE embeddings. **c**, Same as **b** for UMAP embeddings computed with various values of n neighbors (called nn in the legend).

**S4:**
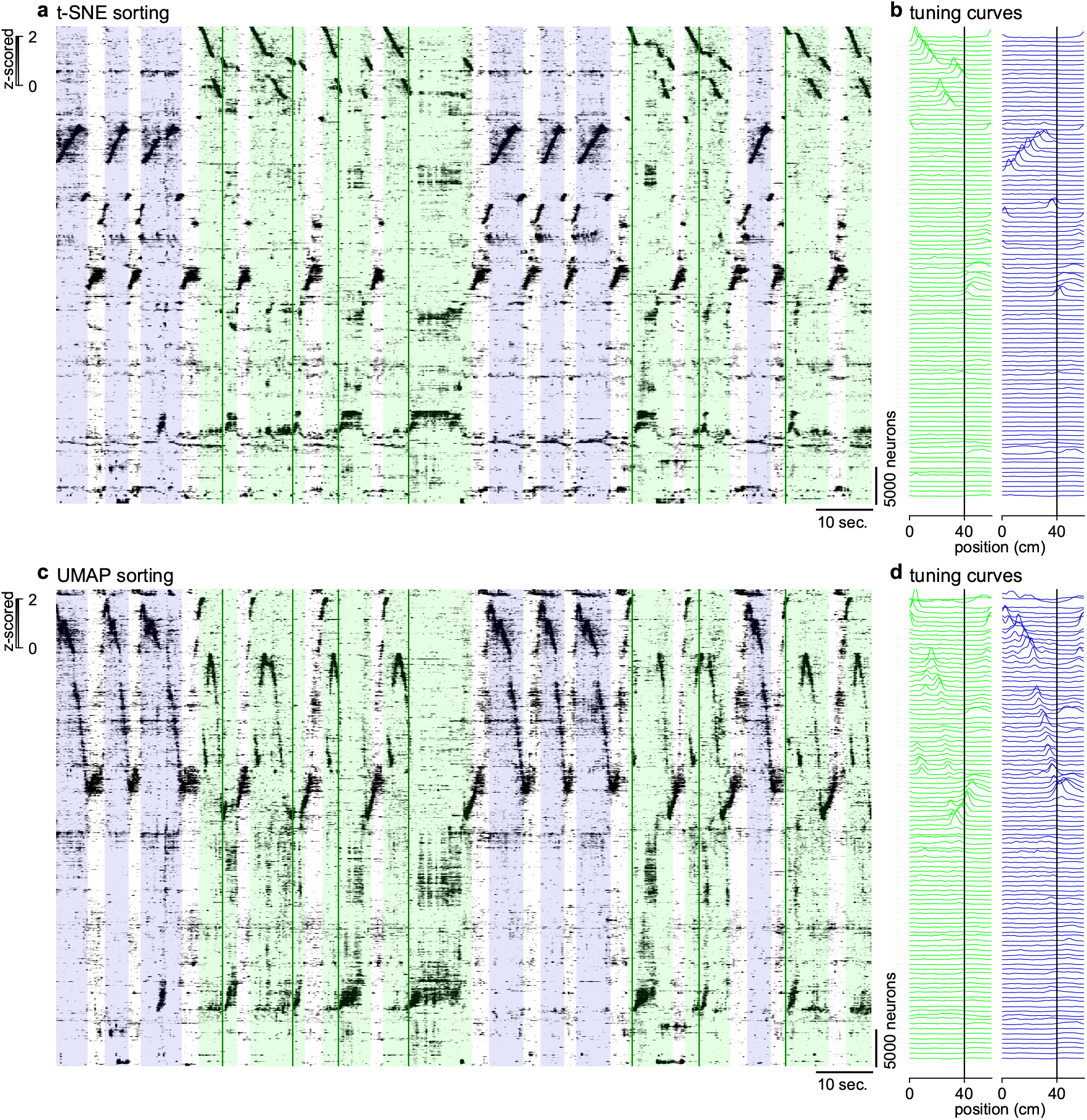
Neural activity from a virtual reality task sorted by t-SNE and UMAP. **a-b**, Same as Figure3d-e using sorting from the t-SNE algorithm. **c-d**, Same as Figure3d-e using sorting from the UMAP algorithm.

**S5:**
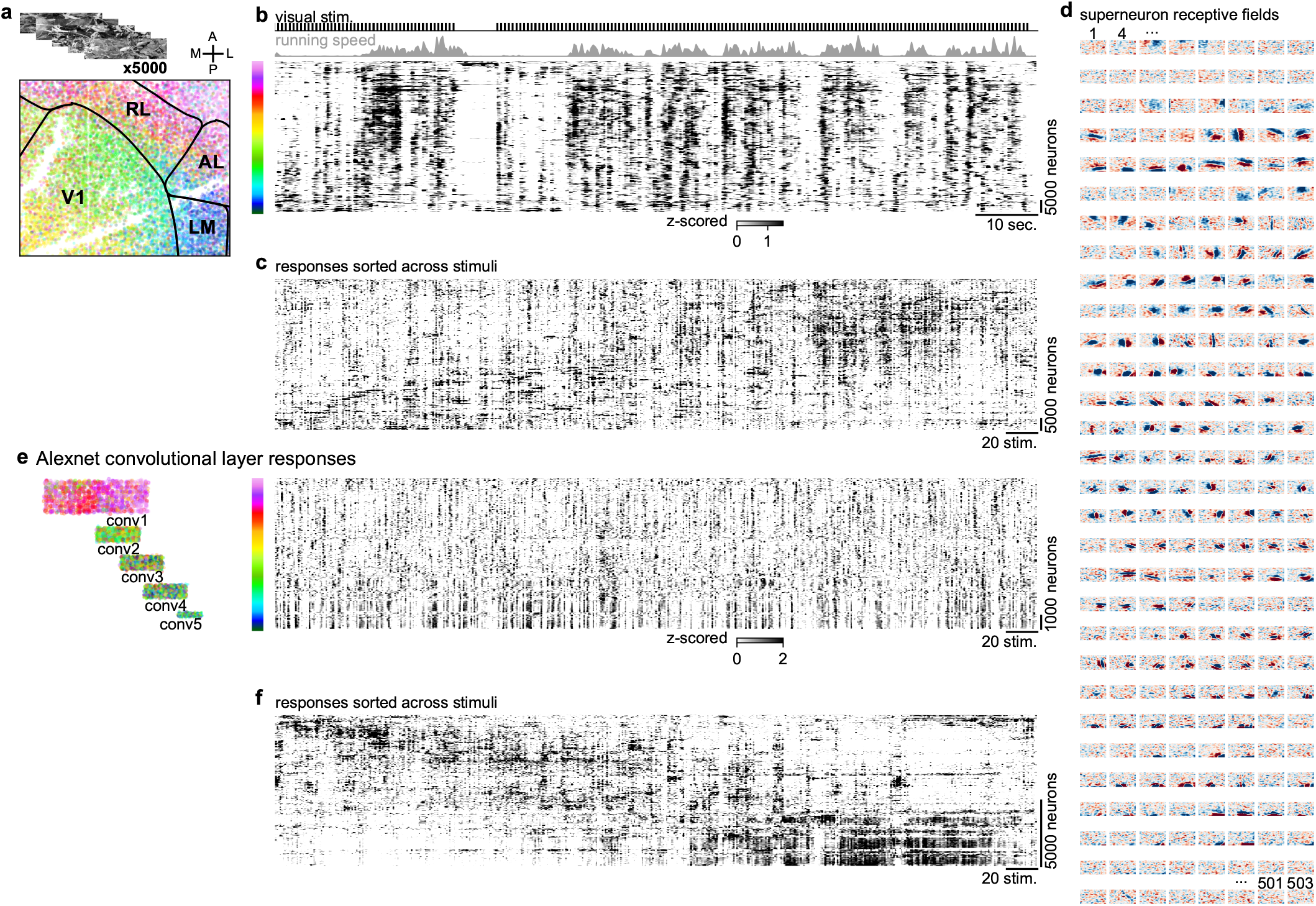
Responses of real and artificial neurons to natural images. **a**, 5,000 natural images were shown to a mouse during a two-photon calcium imaging recording from V1 and higher visual areas. **b**, Activity from 69,957 neurons sorted by Rastermap with splitting, and binned into superneurons, plotted with the mouse’s running speed and the visual stimulus times. **c**, Superneuron responses sorted using Rastermap along the stimulus axis, every tenth stimulus is shown out of 5,000. **d**, Linear receptive fields for the superneurons in **b** in the same order. **e**, Alexnet convolutional layer responses to the same 5,000 natural images sorted by Rastermap with splitting. Left: Units in the convolutional layers colored by the Rastermap sorting. Right: Unit activations sorted and binned into superneurons shown across stimuli. **f**, Superneuron responses sorted using Rastermap along the stimulus axis, every tenth stimulus is shown out of 5,000.

**S6:**
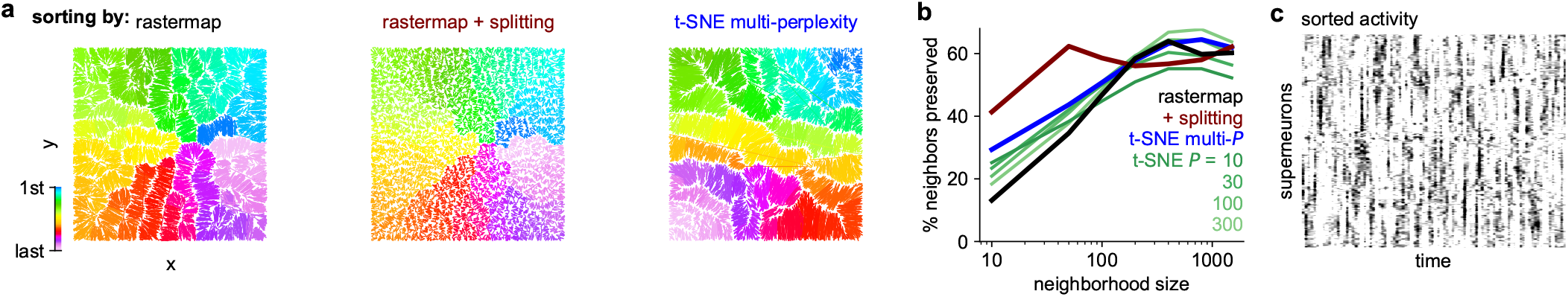
Simulation of neural activity with intrinsic dimensionality of 2. We simulated neural activity with an intrinsic dimensionality of 2 by randomly choosing an *x* and *y* value for each neuron in the range of 0 to 1 and modelling its activity as a place field. **a**, Left: Simulated neurons are plotted at their ground-truth (*x, y*) positions and colored by their position in the Rastermap sorting run with *N*_clusters_ = 100. Middle: Same as the left panel, using Rastermap with splitting, resulting in *N*_clusters_ = 800. Right: Same as the left panel, using t-SNE with multiple perplexities (*P* = (10, 100)) to sort the neurons. **b**, The KNN score for benchmarking embedding algorithms from [25]: the percentage of *k*-nearest neighbors in the original space that are preserved as *k*-nearest neighbors in the embedding space. Shown for Rastermap, Rastermap with splitting, and t-SNE with various perplexities. **c**, Simulated activity sorted by Rastermap with splitting.

## References

[1] Na Ji, Jeremy Freeman, and Spencer L Smith. Technologies for imaging neural activity in large volumes. Nature neuroscience, 19(9):1154–1164, 2016.

[2] Nicholas A Steinmetz, Christof Koch, Kenneth D Harris, and Matteo Carandini. Challenges and opportunities for large-scale electrophysiology with neuropixels probes. Current opinion in neurobiology, 50:92–100, 2018.

[3] Alison I Weber and Jonathan W Pillow. Capturing the dynamical repertoire of single neurons with generalized linear models. Neural computation, 29(12):3260–3289, 2017.

[4] Panayiota Poirazi and Athanasia Papoutsi. Illuminating dendritic function with computational models. Nature Reviews Neuroscience, 21(6):303–321, 2020.

[5] John P Cunningham and Byron M Yu. Dimensionality reduction for large-scale neural recordings. Nature Neuroscience, 17(11):1500, 2014.

[6] Rich Pang, Benjamin J. Lansdell, and Adrienne L. Fairhall. Dimensionality reduction in neuroscience. Current Biology, 26(14):R656–R660, 2016.

[7] Tatiana A. Engel, Nicholas A. Steinmetz, Marc A. Gieselmann, Alexander Thiele, Tirin Moore, and Kwabena Boahen. Selective modulation of cortical state during spatial attention. Science, 354(6316):1140–1144, 2016.

[8] Bingni W Brunton, Lise A Johnson, Jeffrey G Ojemann, and J Nathan Kutz. Extracting spatial– temporal coherent patterns in large-scale neural recordings using dynamic mode decomposition. Journal of neuroscience methods, 258:1–15, 2016.

[9] A. Aldo Faisal, Luc P. J. Selen, and Daniel M. Wolpert. Noise in the nervous system. Nature reviews. Neuroscience, 9(4):292–303, Apr 2008. 18319728[pmid].

[10] G Bard Ermentrout, Roberto F Galán, and Nathaniel N Urban. Reliability, synchrony and noise. Trends in neurosciences, 31(8):428–434, 2008.

[11] Marlene R Cohen and Adam Kohn. Measuring and interpreting neuronal correlations. Nature Neuroscience, 14(7):811–819, 2011.

[12] I-Chun Lin, Michael Okun, Matteo Carandini, and Kenneth D Harris. The nature of shared cortical variability. Neuron, 87(3):644–656, 2015.

[13] Katie A Ferguson and Jessica A Cardin. Mechanisms underlying gain modulation in the cortex. Nature Reviews Neuroscience, 21(2):80–92, 2020.

[14] Lilach Avitan, Zac Pujic, Jan Mö lter, Matthew Van De Poll, Biao Sun, Haotian Teng, Rumelo Amor, Ethan K. Scott, and Geoffrey J. Goodhill. Spontaneous activity in the zebrafish tectum reorganizes over development and is influenced by visual experience. Current Biology, 27(16):2407 –2419, 2017.

[15] Carsen Stringer, Marius Pachitariu, Nicholas Steinmetz, Charu Bai Reddy, Matteo Carandini, and Kenneth D Harris. Spontaneous behaviors drive multidimensional, brainwide activity. Science, 364(6437):eaav7893, 2019.

[16] Carsen Stringer, Marius Pachitariu, Nicholas Steinmetz, Matteo Carandini, and Kenneth D Harris. High-dimensional geometry of population responses in visual cortex. Nature, 571(7765):361–365, 2019.

[17] Frederic Lanore, N Alex Cayco-Gajic, Harsha Gurnani, Diccon Coyle, and R Angus Silver. Cerebellar granule cell axons support highdimensional representations. Nature Neuroscience, 24(8):1142–1150, 2021.

[18] Laurens van der Maaten and Geoffrey Hinton. Visualizing data using t-sne. Journal of machine learning research, 9(Nov):2579–2605, 2008.

[19] Leland McInnes, John Healy, and James Melville. Umap: Uniform manifold approximation and projection for dimension reduction. arXiv preprint arXiv:1802.03426, 2018.

[20] Xiuye Chen, Yu Mu, Yu Hu, Aaron T Kuan, Maxim Nikitchenko, Owen Randlett, Alex B Chen, Jeffery P Gavornik, Haim Sompolinsky, Florian Engert, and Misha B Ahrens. Brain-wide organization of neuronal activity and convergent sensorimotor transformations in larval zebrafish. Neuron, 100(4):876–890, 2018.

[21] Dmitry Kobak and George C Linderman. Initialization is critical for preserving global data structure in both t-sne and umap. Nature biotechnology, 39(2):156–157, 2021.

[22] Tara Chari, Joeyta Banerjee, and Lior Pachter. The specious art of single-cell genomics. BioRxiv, pages 2021–08, 2021.

[23] Artur Luczak, Peter Barthó, and Kenneth D Haris. Spontaneous events outline the realm of possible sensory responses in neocortical populations. Neuron, 62(3):413–425, 2009.

[24] Karunesh Ganguly, Lavi Secundo, Gireeja Ranade, Amy Orsborn, Edward F. Chang, Dragan F. Dimitrov, Jonathan D. Wallis, Nicholas M. Barbaro, Robert T. Knight, and Jose M. Carmena. Cortical representation of ipsilateral arm movements in monkey and man. Journal of Neuroscience, 29(41):12948–12956, 2009.

[25] Dmitry Kobak and Philipp Berens. The art of using t-sne for single-cell transcriptomics. Nature communications, 10(1):5416, 2019.

[26] John A Lee, Diego H Peluffo-Ordóñez, and Michel Verleysen. Multi-scale similarities in stochastic neighbour embedding: Reducing dimensionality while preserving both local and global structure. Neurocomputing, 169:246–261, 2015.

[27] Michael Jü nger, Gerhard Reinelt, and Giovanni Rinaldi. The traveling salesman problem. Handbooks in operations research and management science, 7:225–330, 1995.

[28] Georges A Croes. A method for solving traveling-salesman problems. Operations research, 6(6):791–812, 1958.

[29] Shen Lin. Computer solutions of the traveling salesman problem. Bell System Technical Journal, 44(10):2245–2269, 1965.

[30] Siu Kwan Lam, Antoine Pitrou, and Stanley Seibert. Numba: A llvm-based python jit compiler. In Proceedings of the Second Workshop on the LLVM Compiler Infrastructure in HPC, page 7. ACM, 2015.

[31] Carsen Stringer, Michalis Michaelos, Dmitri Tsyboulski, Sarah E Lindo, and Marius Pachitariu. High-precision coding in visual cortex. Cell, 184(10):2767–2778, 2021.

[32] Nicholas James Sofroniew, Daniel Flickinger, Jonathan King, and Karel Svoboda. A large field of view two-photon mesoscope with subcellular resolution for in vivo imaging. Elife, 5:e14472, 2016.

[33] Dmitri Tsyboulski, Natalia Orlova, Fiona Griffin, Sam Seid, Jerome Lecoq, and Peter Saggau. Remote focusing system for simultaneous dual-plane mesoscopic multiphoton imaging. bioRxiv, page 503052, 2018.

[34] Marius Pachitariu, Carsen Stringer, Sylvia Schrö der, Mario Dipoppa, L Federico Rossi, Matteo Carandini, and Kenneth D Harris. Suite2p: beyond 10,000 neurons with standard two-photon microscopy. bioRxiv, 2016.

[35] Atika Syeda, Lin Zhong, Renee Tung, Will Long, Marius Pachitariu, and Carsen Stringer. Facemap: a framework for modeling neural activity based on orofacial tracking. bioRxiv, 2022.

[36] Cristopher M. Niell and Michael P. Stryker. Modulation of Visual Responses by Behavioral State in Mouse Visual Cortex. Neuron, 65(4):472–479, 2010.

[37] Martin Vinck, Renata Batista-Brito, Ulf Knoblich, and Jessica A. Cardin. Arousal and Locomotion Make Distinct Contributions to Cortical Activity Patterns and Visual Encoding. Neuron, 86(3):740–754, 2015.

[38] Laura N Driscoll, Noah L Pettit, Matthias Minderer, Selmaan N Chettih, and Christopher D Harvey. Dynamic reorganization of neuronal activity patterns in parietal cortex. Cell, 170(5):986–999, 2017.

[39] Janelle MP Pakan, Stephen P Currie, Lukas Fischer, and Nathalie L Rochefort. The impact of visual cues, reward, and motor feedback on the representation of behaviorally relevant spatial locations in primary visual cortex. Cell reports, 24(10):2521–2528, 2018.

[40] Michael Krumin, Julie J Lee, Kenneth D Harris, and Matteo Carandini. Decision and navigation in mouse parietal cortex. elife, 7:e42583, 2018.

[41] Andres D Grosmark and Gyö rgy Buzsáki. Diversity in neural firing dynamics supports both rigid and learned hippocampal sequences. Science, 351(6280):1440–1443, 2016.

[42] Andres D Grosmark, John Long, and Gyö rgy Buzsáki. Recordings from hippocampal area ca1, pre, during and post novel spatial learning. CRCNS. org, 10:K0862DC5, 2016.

[43] Xiuye Chen, Yu Mu, Aaron Kuan, Maxim Nikitchenko, Owen Randlett, Alex Chen, Jeffery Gavornik, Haim Sompolinsky, Florian Engert, and Misha B. Ahrens. Whole-brain light-sheet imaging data. https://janelia.figshare.com/articles/dataset/Wholebrainlight-sheetimagingdata/7272617, 2019.

[44] Simon Musall, Matthew T Kaufman, Ashley L Juavinett, Steven Gluf, and Anne K Churchland. Single-trial neural dynamics are dominated by richly varied movements. Nature neuroscience, 22(10):1677–1686, 2019.

[45] Anne K Churchland, Simon Musall, Matthew T Kaufman, Ashley L Juavinett, and Steven Gluf. Dataset from: Single-trial neural dynamics are dominated by richly varied movements. http://repository.cshl.edu/id/eprint/38599/, 2019.

[46] Claudia E Feierstein, Ruben Portugues, and Michael B Orger. Seeing the whole picture: a comprehensive imaging approach to functional mapping of circuits in behaving zebrafish. Neuroscience, 296:26–38, 2015.

[47] Lilach Avitan and Carsen Stringer. Not so spontaneous: Multi-dimensional representations of behaviors and context in sensory areas. Neuron, 2022.

[48] Chi Ren and Takaki Komiyama. Characterizing cortex-wide dynamics with wide-field calcium imaging. Journal of Neuroscience, 41(19):4160–4168, 2021.

[49] Antonin Raffin, Ashley Hill, Adam Gleave, Anssi Kanervisto, Maximilian Ernestus, and Noah Dormann. Stable-baselines3: Reliable reinforcement learning implementations. Journal of Machine Learning Research, 22(268):1–8, 2021.

[50] Antonin Raffin. Rl baselines3 zoo. https://github.com/DLR-RM/rl-baselines3-zoo, 2020.

[51] Jeremy Freeman, Nikita Vladimirov, Takashi Kawashima, Yu Mu, Nicholas J Sofroniew, Davis V Bennett, Joshua Rosen, Chao-Tsung Yang, Loren L Looger, and Misha B Ahrens. Mapping brain activity at scale with cluster computing. Nature methods, 11(9):941–950, 2014.

[52] Emily L Mackevicius, Andrew H Bahle, Alex H Williams, Shijie Gu, Natalia I Denisenko, Mark S Goldman, and Michale S Fee. Unsupervised discovery of temporal sequences in highdimensional datasets, with applications to neuroscience. Elife, 8:e38471, 2019.

[53] Dmitry Kobak, Wieland Brendel, Christos Constantinidis, Claudia E Feierstein, Adam Kepecs, Zachary F Mainen, Xue-Lian Qi, Ranulfo Romo, Naoshige Uchida, and Christian K Machens. Demixed principal component analysis of neural population data. elife, 5:e10989, 2016.

[54] Chethan Pandarinath, K Cora Ames, Abigail A Russo, Ali Farshchian, Lee E Miller, Eva L Dyer, and Jonathan C Kao. Latent factors and dynamics in motor cortex and their application to brain– machine interfaces. Journal of Neuroscience, 38(44):9390–9401, 2018.

[55] Alex H Williams, Tony Hyun Kim, Forea Wang, Saurabh Vyas, Stephen I Ryu, Krishna V Shenoy, Mark Schnitzer, Tamara G Kolda, and Surya Ganguli. Unsupervised discovery of demixed, low-dimensional neural dynamics across multiple timescales through tensor component analysis. Neuron, 98(6):1099–1115, 2018.

[56] Ding Zhou and Xue-Xin Wei. Learning identifiable and interpretable latent models of highdimensional neural activity using pi-vae. Advances in Neural Information Processing Systems, 33:7234–7247, 2020.

[57] Steffen Schneider, Jin Hwa Lee, and Mackenzie Weygandt Mathis. Learnable latent embeddings for joint behavioural and neural analysis. Nature, pages 1–9, 2023.

[58] Mahathi Ramaswamy, Ruey-Kuang Cheng, and Suresh Jesuthasan. Identification of gabaergic neurons innervating the zebrafish lateral habenula. European Journal of Neuroscience, 52(8):3918–3928, 2020.

[59] Joel Bauer, Simon Weiler, Martin HP Fernholz, David Laubender, Volker Scheuss, Mark Hü bener, Tobias Bonhoeffer, and Tobias Rose. Limited functional convergence of eye-specific inputs in the retinogeniculate pathway of the mouse. Neuron, 109(15):2457–2468, 2021.

[60] Jordan S Farrell, Roberto Colangeli, Ao Dong, Antis G George, Kwaku Addo-Osafo, Philip J Kingsley, Maria Morena, Marshal D Wolff, Barna Dudok, Kaikai He, Toni A. Patrick, Keith A. Sharkey, Sachin Patel, Lawrence J. Marnett, Matthew N. Hill, Yulong Li, G. Campbell Teskey, and Ivan Soltesz. In vivo endocannabinoid dynamics at the timescale of physiological and pathological neural activity. Neuron, 109(15):2398–2403, 2021.

[61] Laurie-Anne Lamiré, Martin Haesemeyer, Florian Engert, Michael Granato, and Owen Randlett. Functional and pharmacological analyses of visual habituation learning in larval zebrafish. bioRxiv, pages 2022–06, 2022.

[62] Seetha Krishnan, Chad Heer, Chery Cherian, and Mark EJ Sheffield. Reward expectation extinction restructures and degrades ca1 spatial maps through loss of a dopaminergic reward proximity signal. Nature Communications, 13(1):6662, 2022.

[63] Helen Farrants, Yichun Shuai, William C Lemon, Christian Monroy Hernandez, Shang Yang, Ronak Patel, Guanda Qiao, Michelle S Frei, Jonathan B Grimm, Timothy L Hanson, Filip Tomaska, Glenn C Turner, Carsen Stringer, Philipp J Keller, Abraham G Beyene, Yao Chen, Yajie Liang, Luke D Lavis, and Eric R Schreiter. A modular chemigenetic calcium indicator enables in vivo functional imaging with near-infrared light. bioRxiv, 2023.

[64] Jie-Yoon Yang, Thomas F O’Connell, Wei-Mien M Hsu, Matthew S Bauer, Kristina V Dylla, Tatyana O Sharpee, and Elizabeth J Hong. Restructuring of olfactory representations in the fly brain around odor relationships in natural sources. bioRxiv, pages 2023–02, 2023.

[65] Célian Bimbard, Timothy PH Sit, Anna Lebedeva, Charu B Reddy, Kenneth D Harris, and Matteo Carandini. Behavioral origin of sound-evoked activity in mouse visual cortex. Nature neuroscience, 26(2):251–258, 2023.

[66] Lauren E Wool, Armin Lak, Matteo Carandini, and Kenneth D Harris. Mouse frontal cortex nonlinearly encodes sensory, choice and outcome signals. bioRxiv, pages 2023–05, 2023.

[67] Camille Testard, Sebastien Tremblay, Felipe Parodi, Ronald W DiTullio, Arianna Acevedo-Ithier, Kristin Gardiner, Konrad Paul Kording, and Michael Platt. Neural signatures of natural behavior in socializing macaques. bioRxiv, pages 2023–07, 2023.

[68] Thomas A Pologruto, Bernardo L Sabatini, and Karel Svoboda. Scanimage: flexible software for operating laser scanning microscopes. Biomedical engineering online, 2(1):13, 2003.

[69] Mario Kleiner, David Brainard, Denis Pelli, Allen Ingling, Richard Murray, Christopher Broussard, et al. What’s new in psychtoolbox-3. Perception, 36(14):1, 2007.

[70] Maximilian Joesch and Markus Meister. A neuronal circuit for colour vision based on rod–cone opponency. Nature, 532(7598):236–239, 2016.

[71] Nader Nikbakht and Mathew E Diamond. Conserved visual capacity of rats under red light. eLife, 10:e66429, jul 2021.

[72] Johannes Friedrich, Pengcheng Zhou, and Liam Paninski. Fast online deconvolution of calcium imaging data. PLOS Computational Biology, 13(3):e1005423, 2017.

[73] Marius Pachitariu, Carsen Stringer, and Kenneth D Harris. Robustness of spike deconvolution for neuronal calcium imaging. Journal of Neuroscience, 38(37):7976–7985, 2018.

[74] Stefan Van Der Walt, S Chris Colbert, and Gael Varoquaux. The numpy array: a structure for efficient numerical computation. Computing in Science & Engineering, 13(2):22, 2011.

[75] Eric Jones, Travis Oliphant, Pearu Peterson, et al. SciPy: Open source scientific tools for Python, 2001.

[76] Fabian Pedregosa, Gaë l Varoquaux, Alexandre Gramfort, Vincent Michel, Bertrand Thirion, Olivier Grisel, Mathieu Blondel, Peter Prettenhofer, Ron Weiss, Vincent Dubourg, et al. Scikitlearn: Machine learning in python. Journal of machine learning research, 12(Oct):2825–2830, 2011.

[77] PyQT. Pyqt reference guide. 2012.

[78] Luke Campagnola. Scientific graphics and gui library for python. https://github.com/pyqtgraph/pyqtgraph, 2020.

[79] Thomas Kluyver, Benjamin Ragan-Kelley, Fernando Pérez, Brian E Granger, Matthias Bussonnier, Jonathan Frederic, Kyle Kelley, Jessica B Hamrick, Jason Grout, Sylvain Corlay, et al. Jupyter notebooks-a publishing format for reproducible computational workflows. In ELPUB, pages 87–90, 2016.

[80] John D Hunter. Matplotlib: A 2d graphics environment. Computing in science & engineering, 9(3):90, 2007.

[81] Eric W Weisstein. Peano curve. https://mathworld.wolfram.com/, 2015.

[82] Joshua B Tenenbaum, Vin de Silva, and John C Langford. A global geometric framework for nonlinear dimensionality reduction. science, 290(5500):2319–2323, 2000.

[83] Mikhail Belkin and Partha Niyogi. Laplacian eigenmaps for dimensionality reduction and data representation. Neural computation, 15(6):1373–1396, 2003.

[84] Pavlin G. Poličar, Martin Stražar, and Blaž Zupan. opentsne: a modular python library for t-sne dimensionality reduction and embedding. bioRxiv, 2019.

[85] Will Dabney, Mark Rowland, Marc Bellemare, and Rémi Munos. Distributional reinforcement learning with quantile regression. Proceedings of the AAAI Conference on Artificial Intelligence, 32(1), Apr. 2018.

[86] Volodymyr Mnih, Koray Kavukcuoglu, David Silver, Andrei A Rusu, Joel Veness, Marc G Bellemare, Alex Graves, Martin Riedmiller, Andreas K Fidjeland, Georg Ostrovski, et al. Human-level control through deep reinforcement learning. nature, 518(7540):529–533, 2015.

[87] Alex Krizhevsky, Ilya Sutskever, and Geoffrey E Hinton. Imagenet classification with deep convolutional neural networks. Advances in neural information processing systems, 25, 2012.

[88] Jia Deng, Wei Dong, Richard Socher, Li-Jia Li, Kai Li, and Li Fei-Fei. Imagenet: A large-scale hierarchical image database. In IEEE Conference on Computer Vision and Pattern Recognition., pages 248–255. IEEE, 2009.

